# Targeting of Cdc42 GTPase in regulatory T cells unleashes anti-tumor T cell immunity

**DOI:** 10.1101/2021.09.23.461402

**Authors:** Khalid W Kalim, Jun-Qi Yang, Mark Wunderlich, Vishnu Modur, Phuong Nguyen, Yuan Li, Ting Wen, Ashley Kuenzi Davis, Ravinder Verma, Q. Richard Lu, Anil G Jegga, Yi Zheng, Fukun Guo

## Abstract

Regulatory T (Treg) cells play an important role in maintaining immune tolerance through inhibiting effector T cell function. In the tumor microenvironment, Treg cells are utilized by tumor cells to counteract effector T cell-mediated tumor killing. Targeting Treg cells may thus unleash the anti-tumor activity of effector T cells. While systemic depletion of Treg cells can cause excessive effector T cell responses and subsequent autoimmune diseases, controlled targeting of Treg cells may benefit cancer patients. Here we show that Treg cell-specific heterozygous deletion or pharmacological targeting of Cdc42 GTPase does not affect Treg cell numbers but induces Treg cell plasticity, leading to anti-tumor T cell immunity without detectable autoimmune reactions. Cdc42 targeting potentiates an immune checkpoint blocker anti-PD-1 antibody-mediated T cell response against mouse and human tumors. Mechanistically, Cdc42 targeting induces Treg cell plasticity and unleashes antitumor T cell immunity through carbonic anhydrase I-mediated pH changes. Thus, rational targeting of Cdc42 in Treg cells holds therapeutic promises in cancer immunotherapy.

**Significance:** Effector T lymphocytes promote autoimmune diseases but have potential to kill tumor cells. However, cancer cells can evade T cell-mediated killing in part by utilizing regulatory T (Treg) cells to inhibit effector T cell function. Here we show that Treg cell-specific heterozygous deletion of Cdc42 gene that encodes Cdc42 GTPase dampens Treg cell fitness through carbonic anhydrase I-mediated pH changes, leading to anti-tumor T cell immunity. Pharmacological targeting of Cdc42 mimics genetic deletion of Cdc42 in impairing Treg cell fitness and evoking anti-tumor T cell immunity. Importantly, Cdc42 targeting does not appear to cause systemic autoimmunity. Given that current cancer immunotherapies only demonstrate limited clinical efficacies, our findings may open a new avenue for cancer immunotherapy.

## Introduction

CD4^+^ (T helper 1 (Th1), Th2, Th17) and CD8^+^ effector T lymphocytes have the potential to kill tumor cells. However, tumor cells can evade immune surveillance partly by engaging immune checkpoint proteins (e.g. PD-1, CTLA-4) on effector T cells to cause their exhaustion (1). Several immune checkpoint inhibitors (ICIs) (e.g. Anti-PD-1) have been approved by FDA for the clinical treatment of a number of cancer types (2). However, these immune checkpoint inhibitors only demonstrate clinical efficacy in a small proportion of cancer patients (3), necessitating improved cancer immunotherapies.

CD4^+^Foxp3^+^ regulatory T (Treg) cells are important for maintaining immune tolerance, primarily by inhibiting effector T cells (4). Consequently, defects in Treg cell homeostasis or their suppressive function can lead to excessive effector T cell responses and autoimmunity (5, 6). In the tumor microenvironment (TME), Treg cells contribute to tumor immune evasion (7, 8). Depletion of Treg cells may thus benefit cancer patients (9). However, systemic removal of Treg cells likely causes autoimmune diseases (9). It has recently been suggested that cancer may be treated by induction of Treg cell plasticity that in general is reflected by loss of stable expression of Foxp3, the signature transcription factor and an important functional marker of Treg cells, effector T cell reprogramming, and/or impairment of Treg cell function (10–13). The mechanisms underlying Treg cell plasticity remain poorly defined, understanding of which may provide novel biological targets for cancer immunotherapy.

Cdc42 is a Rho family GTPase that regulates a variety of cellular events including cell proliferation, survival, actin cytoskeletal organization, and migration (14). We have recently reported that Treg cell-specific homozygous gene deletion of *Cdc42* decreases Treg cell numbers, induces Treg cell plasticity, and causes early, fatal inflammatory diseases (15). This suggests that Cdc42 is important for maintenance of Treg cell homeostasis and fitness and for controlling autoimmunity. Here we report that Treg cell-specific heterozygous gene deletion of *Cdc42* also induces Treg cell plasticity but does not reduce Treg cell number nor does it cause autoimmunity. Importantly, heterozygous deletion of *Cdc42* activates anti-tumor T cell immunity. Mechanistically, heterozygous deletion of *Cdc42* induces Treg cell plasticity and unleashes anti-tumor T cell immunity through upregulation of carbonic anhydrase I (CAI) that functions to modulate cellular pH by catalyzing the hydration of CO_2_ to HCO_3_^-^ and H^+^ (16). A Cdc42 inhibitor, CASIN (17), mimics heterozygous deletion of *Cdc42* in inducing Treg cell plasticity and causes anti-tumor T cell immunity, without incurring autoimmune responses. Additionally, CASIN synergizes with Anti-PD-1 in triggering anti-tumor T cell immunity. Our findings indicate that pharmacological titration of Cdc42 to induce Treg cell plasticity without altering their homeostasis may be useful for immunotherapy modulations without causing autoimmunity.

## Results

### Heterozygous Deletion of *Cdc42* Induces Treg Cell Plasticity but Does Not Cause Autoimmunity

By crossing *Cdc42^Flox/Flox^* mice with *Foxp3^YFP-Cre^* mice, we generated *Cdc42^Flox/+^Foxp3^YFP-Cre^* mice that harbor heterozygous knockout of *Cdc42* specifically in Treg cells. We found that heterozygous knockout of *Cdc42* did not affect Treg cell homeostasis (Fig. 1*A*), proliferation (Fig. 1*B*), and survival (Fig. 1*C*). A variety of Treg cell functional markers such as PD-1, CTLA-4, ICOS, GITR, CD39, and CD73 also remained unchanged (Fig. 1*D*). *Cdc42^Flox/+^Foxp3^YFP-Cre^* mice had no visible inflammatory disorders. H&E staining showed no inflammatory cell infiltration into the colon, liver, lung, and kidney of *Cdc42^Flox/+^Foxp3^YFP-Cre^* mice (Fig. 1*E*). However, *Cdc42^Flox/+^Foxp3^YFP-Cre^* Treg cells showed reduced expression of Foxp3 (Fig. 1*F*) and increased methylation in conserved noncoding sequence 2 (CNS2) of the *Foxp3* locus (Fig. 1*G*), a CpG-rich Foxp3 intronic cis-element whose hypomethylation is important for stable Foxp3 expression (13). The increased methylation in CNS2 of the Foxp3 locus was associated with increased expression of DNA methyltransferase 3A (DNMT3a) *(SI Appendix* Fig. S1). *Cdc42^Flox/+^Foxp3^YFP-Cre^* Treg cells underwent effector T cell reprogramming, as evidenced by increased expression of Th2 cytokine IL-4 and Th1 cytokine IFN-γ but not Th17 cytokine IL-17 (Fig. 1*H* and *SI Appendix* Fig. S2*A*). Although IFN-γ-expressing CD8^+^ effector T cells were not altered (Fig. 1*I* and *SI Appendix* Fig. S2*B*), IL-4- and IFN-γ-expressing CD4^+^ effector T cells (CD4^+^Foxp3^-^) were increased in *Cdc42^Flox/+^Foxp3^YFP-Cre^* mice (Fig. 1*J* and *SI Appendix* Fig. S2*A*). *Ex vivo* culture of Treg cells found that heterozygous loss of *Cdc42* caused more Treg cell conversion to CD4^+^Foxp3^-^ ex-Treg cells (Fig. 1*K*). To determine whether the same occurs *in vivo*, we took advantage of *Foxp3*^eGFP-Cre-ERT2^*Rosa26^eYFP^* mice. In *Foxp3*^eGFP-Cre-ERT2^*Rosa26^eYFP^* mice, treatment with tamoxifen induces the expression of Cre and GFP in Foxp3^+^ cells. Cre expression causes excision of floxed stop codon in the *Rosa26* locus, leading to YFP expression. Thus, Treg cells from *Foxp3*^eGFP-Cre-ERT2^*Rosa26^eYFP^* mice are marked by both GFP and YFP. When Treg cells convert to CD4^+^Foxp3^-^ ex-Treg cells, their GFP is lost but YFP is retained, because the Rosa26 locus is ubiquitously expressed. We crossed *Foxp3*^eGFP-Cre-ERT2^*Rosa26^eYFP^* mice with *Cdc42^Flox/Flox^* mice and found that the resultant *Cdc42^Flox/+^Foxp3*^eGFP-Cre-ERT2^*Rosa26^eYFP^* mice contained more GFP^-^YFP^+^ ex-Treg cells, compared to *Cdc42*^+/+^*Foxp3*^eGFP-Cre-ERT2^*Rosa26^eYFP^* mice (Fig. 1*L*). Together, these results indicate that heterozygous loss of *Cdc42* primes Treg cell plasticity without invoking autoimmune responses.

**Fig. 1.**
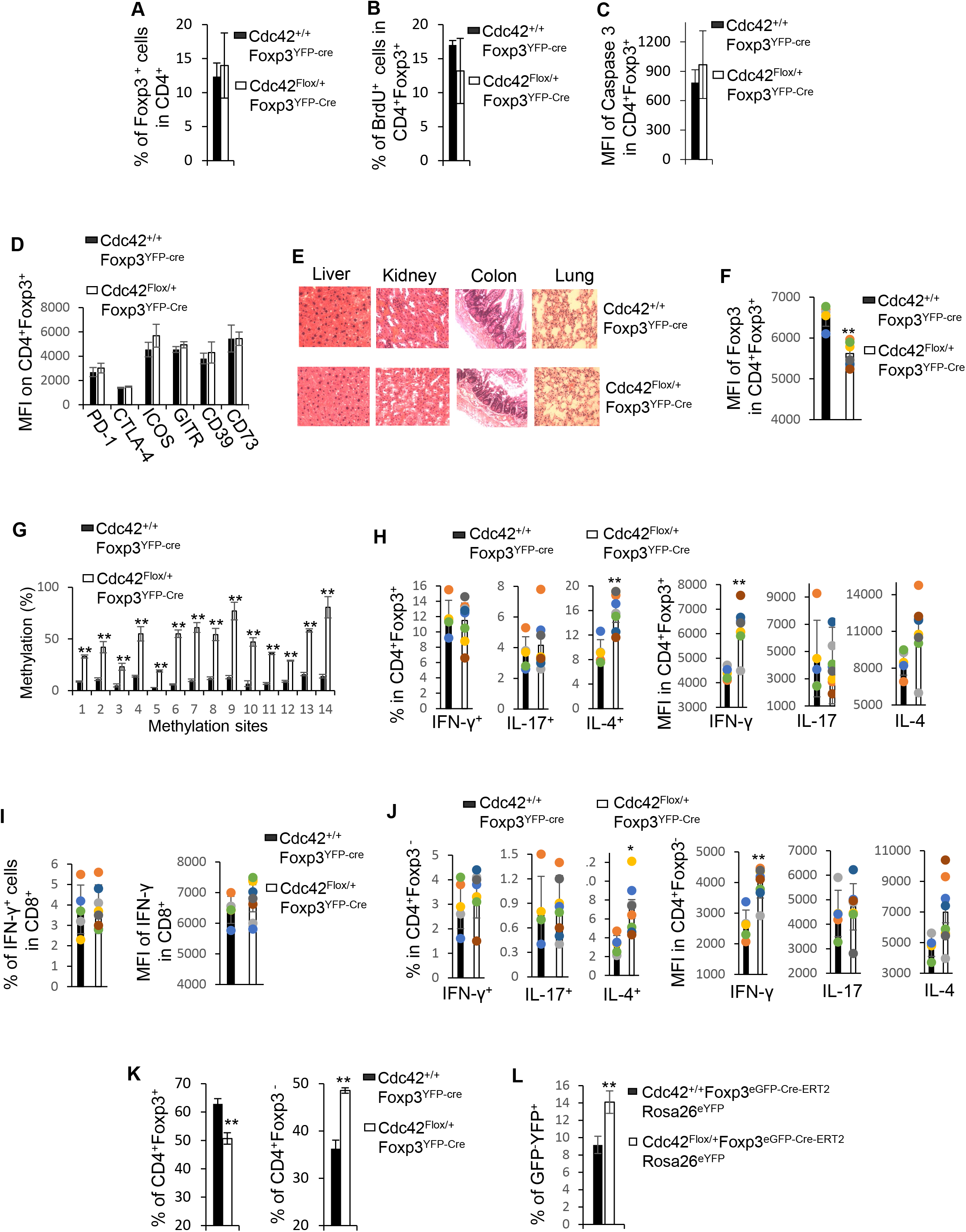
Heterozygous loss of *Cdc42* in Treg cells induces Treg cell plasticity and increases CD4^+^ effector T cells but does not result in autoimmunity. (*A*) Flow cytometry analysis of the percentages of splenic Treg cells in wild type (*Cdc42^+/+^Foxp3^YFP-Cre^*) and *Cdc42* heterozygous knockout (*Cdc42^Flox/+^Foxp3^YFP-Cre^*) mice. (*B*) Flow cytometry analysis of Treg cell proliferation (Brdu incorporation). (*C*) Flow cytometry analysis of Treg cell apoptosis (expression of cleaved caspase 3). (*D*) Flow cytometry analysis of the expression of Treg cell functional markers. (*E*) H&E staining of the indicated organs. (*F*) Flow cytometry analysis of the expression of Foxp3 in Treg cells. (*G*) Bisulfite pyrosequencing analysis of the methylation status of 14 methylation sites of the Foxp3 CNS2 in Treg cells. Data are expressed as percentage of methylation which indicates percent cells with a particular methylation site methylated. *(H-J)* Flow cytometry analysis of the expression (percentages and MFI) of IFN-γ, IL-17 and/or IL-4 in Treg cells (*H*), CD8^+^ T cells (*I*) and CD4^+^ T cells (*J*). *(K)* Flow cytometry analysis of induction of ex-Treg cells (CD4^+^Foxp3^-^) *ex vivo.* Purified Treg cells were cultured for 3 days. *(L)* Flow cytometry analysis of ex-Treg cells (GFP^-^YFP^+^) *in vivo. Cdc42^Flox/+^Foxp3*^eGFP-Cre-ERT2^*Rosa26^eYFP^* and *Cdc42^+/+^Foxp3*^eGFP-Cre-ERT2^*Rosa26^eYFP^* mice were treated (i.p.) with 2 mg of tamoxifen daily for 5 consecutive days. Splenic ex-Treg cells were analyzed 6 weeks after tamoxifen injection. (*A-D*), (*F*), and *(H-J)* Error bars indicate SD. n = 5 for *Cdc42*^+/+^*Foxp3^YFP-Cre^* and n = 8 for *Cdc42^Flox/+^Foxp3^YFP-Cre^*. Data are representative of two independent experiments. (*L*) Error bars indicate SD. n = 5. Data are representative of two independent experiments. (*G*) and (*K*) Data are from one experiment with four mice pooled. Error bars indicate SD of triplicates. *p < 0.05; **p < 0.01. MFI: Mean fluorescence intensity.

### Treg Cell-specific Heterozygous Deletion of *Cdc42* Inhibits Tumor Growth

To determine whether *Cdc42^Flox/+^Foxp3^YFP-Cre^* Treg cells can be harnessed for tumor control, we inoculated *Cdc42^Flox/+^Foxp3^YFP-Cre^* and control *Cdc42^+/+^Foxp3^YFP-Cre^* mice with MC38 mouse colon cancer cells. We found that tumor growth (measured by tumor volume) was markedly suppressed in *Cdc42^Flox/+^Foxp3^YFP-Cre^* mice (Fig. 2*A*). *Cdc42^Flox/+^Foxp3^YFP-Cre^* mice showed increased tumor-infiltrating IFN-γ^+^ Treg cells (Fig. 2*B* and *SI Appendix* Fig. S3*A*) and effector T cells (Fig. 2 *C* and *D* and *SI Appendix* Fig. S3 *B* and *C*). The changes in tumor-infiltrating Treg and effector T cells were recapitulated in splenic Treg and effector T cells (*SI Appendix* Fig. S3 *D-F*), suggesting that *Cdc42* heterozygosity generates a global immuno-effect in tumor-bearing mice. Furthermore, heterozygous loss of *Cdc42* in Treg cells inhibited tumor growth of KPC pancreatic cancer cells (Fig. 2*E*). These results suggest that plastic *Cdc42^Flox/+^Foxp3^YFP-Cre^* Treg cells trigger T cell immunity against tumor growth.

**Fig. 2.**
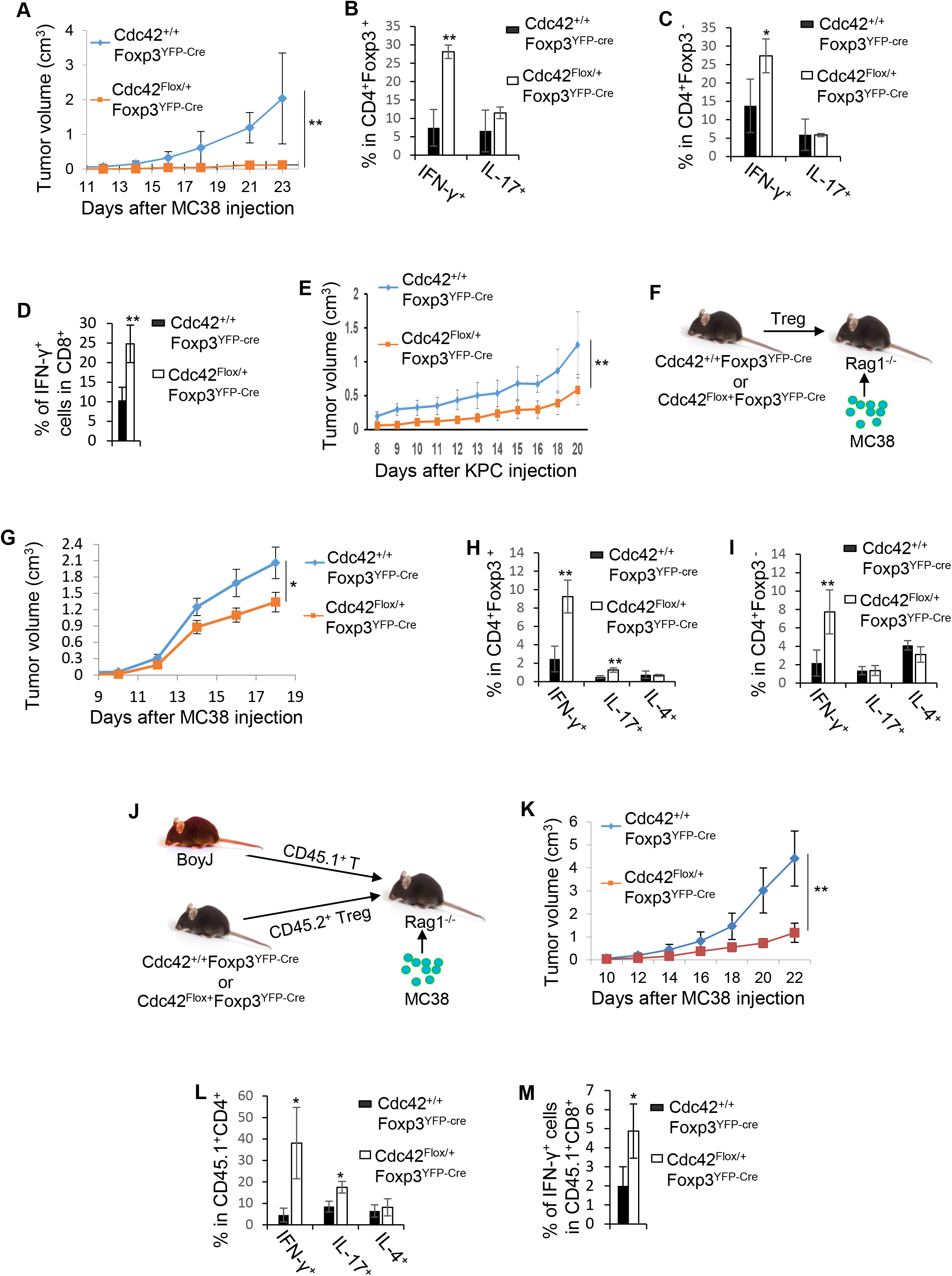
Heterozygous loss of Cdc42 in Treg cells inhibits tumor growth by promoting Treg cell plasticity. (*A*) Tumor growth of MC38 mouse colon cancer cells. (*B-D*) Flow cytometry analysis of the expression of IFN-γ and/or IL-17 in MC38 tumor-infiltrating Treg (CD4^+^Foxp3^+^) (*B*), CD4^+^ effector T (CD4^+^Foxp3^-^) (*C*) and CD8^+^ effector T cells (*D*). (*E*) Tumor growth of KPC mouse pancreatic cancer cells. (*F*) Schematic overview of adoptive Treg cell transfer. (*G*) MC38 Tumor growth in the recipient mice transferred with Treg cells. (*H* and *I*) Flow cytometry analysis of the expression of IFN-γ, IL-17 or IL-4 in MC38 tumor-infiltrating Treg (CD4^+^Foxp3^+^) (*H*) and CD4^+^ effector T cells (ex-Treg, CD4^+^Foxp3^-^) (*I*) in the recipient mice transferred with Treg cells. (*J*) Schematic overview of adoptive co-transfer of Treg (CD45.2^+^) and congenically marked (CD45.1^+^) effector T cells. *(K)* MC38 Tumor growth in the recipient mice transferred with Treg and effector T cells. (*L* and *M*) Flow cytometry analysis of the expression of IFN-γ, IL-17 or IL-4 in MC38 tumor-infiltrating CD45.1^+^CD4^+^ (*L*) and CD45.1^+^CD8^+^ (*M*) effector T cells in the recipient mice transferred with Treg and effector T cells. Error bars indicate SD. n = 6. Data are representative of two independent experiments. *p < 0.05; **p < 0.01.

The increased CD4^+^ effector T cells in tumor-bearing *Cdc42^Flox/+^Foxp3^YFP-Cre^* mice may come from a conversion of plastic Treg cells to effector T cells. To test this possibility, we transferred *Cdc42^Flox/+^Foxp3^YFP-Cre^* and *Cdc42^+/+^Foxp3^YFP-Cre^* Treg cells into immunodeficient *RAG1^-/-^* mice followed by MC38 tumor cell inoculation (Fig. 2*F*). We found that tumor growth was modestly but significantly inhibited in the mice receiving *Cdc42^Flox/+^Foxp3^YFP-Cre^* Treg cells (Fig. 2*G*). In line with this, tumors in the mice receiving *Cdc42^Flox/+^Foxp3^YFP-Cre^* Treg cells showed an increase in plastic Treg cells and in CD4^+^ effector T cells (Fig. 2 *H* and *I*). Hence, plastic *Cdc42^Flox/+^Foxp3^YFP-Cre^* Treg cells indeed converted to CD4^+^ effector T cells (similar to that in *ex vivo* culture (Fig. 1*K*), which may have contributed to the tumor suppression. However, we cannot rule out the possibility that effector T cell activities (e.g. IFN-γ production) in plastic Treg cells may also play a role in the tumor suppression. These data demonstrate that the plasticity of *Cdc42^Flox/+^Foxp3^YFP-Cre^* Treg cells is sufficient for tumor killing.

Plastic Treg cells might be dampened in their suppressive function (11, 13). Thus, the increased CD4^+^ effector T cells in tumor-bearing *Cdc42^Flox/+^Foxp3^YFP-Cre^* mice may not only result from plastic Treg cell conversion to effector T cells, but also compromised function of *Cdc42^Flox/+^Foxp3^YFP-Cre^* Treg cells, which is supported by the increased CD8^+^ effector T cells. To test this possibility, we co-transferred *Cdc42^Flox/+^Foxp3^YFP-Cre^* or *Cdc42^+/+^Foxp3^YFP-Cre^* Treg cells with congenically marked (CD45.1^+^) effector T cells into *Rag1^-/-^* mice followed by tumor cell inoculation (Fig. 2*J*). As expected, tumor growth in the mice receiving *Cdc42^Flox/+^Foxp3^YFP-Cre^* Treg cells and effector T cells was inhibited (Fig. 2*K*). The mice contained increased tumor-infiltrating CD45.1^+^ effector T cells (Fig. 2 *L* and *M*), suggesting that *Cdc42^Flox/+^Foxp3^YFP-Cre^* Treg cells were indeed impaired in their suppressive function. In this setting, the co-transfer of *Cdc42^Flox/+^Foxp3^YFP-Cre^* Treg cells with effector T cells appeared to diminish tumor growth to a greater extent than the transfer of *Cdc42^Flox/+^Foxp3^YFP-Cre^* Treg cells only (Fig. 2*K*, compared to Fig. 2*G*), suggesting that the dampened suppressive function of *Cdc42^Flox/+^Foxp3^YFP-Cre^* Treg cells also contributes to tumor killing in *Cdc42^Flox/+^Foxp3^YFP-Cre^* mice.

Of note, plastic Treg cells (*SI Appendix* Fig. S4 *A* and *B*) and/or the resultant effector T cells (*SI Appendix* Fig. S4 *C* and *D*) in 12-18 months old *Cdc42^Flox/+^Foxp3^YFP-Cre^* mice did not cause spontaneous systemic inflammatory disorders (*SI Appendix* Fig. S4 *E*), similar to that in 6-8 weeks old *Cdc42^Flox/+^Foxp3^YFP-Cre^* mice (Fig. 1) but distinct from that in aged *CNS2^-/-^* mice that show spontaneous lymphoproliferative disease (18). We speculate that the plasticity phenotypes of *Cdc42^Flox/+^Foxp3^YFP-Cre^* Treg cells may be milder than that of *CNS2^-/-^* Treg cells. However, plastic *Cdc42^Flox/+^Foxp3^YFP-Cre^* Treg cells were able to cause anti-tumor T cell immunity in 12-18 months old *Cdc42^Flox/+^Foxp3^YFP-Cre^* mice (*SI Appendix* Fig. S4 *F-I*) as in 6-8 weeks old *Cdc42^Flox/+^Foxp3^YFP-Cre^* mice (Fig. 2).

### Treg Cell-specific Heterozygous Deletion of *Cdc42* Induces Treg Cell Plasticity and Elicits Anti-tumor T Cell Immunity through a Wiskott–Aldrich Syndrome Protein (WASP)-GATA Binding Protein 3 (GATA3)-CAI Signaling Node

To determine the mechanism of heterozygous *Cdc42* loss-induced Treg cell plasticity, we carried out global gene expression profiling of *Cdc42^Flox/+^Foxp3^YFP-Cre^* Treg and *Cdc42^+/+^Foxp3^YFP-Cre^* Treg cells, by RNA-seq (NCBI GEO accession no. GSE181779). Among the gene changes in *Cdc42^Flox/+^Foxp3^YFP-Cre^* Treg cells, *Car1* that encodes CAI was the second most upregulated gene (*SI Appendix* Fig. S5*A*). The expression level of CAI mRNA in *Cdc42^Flox/+^Foxp3^YFP-Cre^* Treg cells was 369 folds more than that in *Cdc42^+/+^Foxp3^YFP-Cre^* Treg cells (*SI Appendix* Fig. S5*B*). Of note, *Car1* was the most upregulated gene in *Cdc42^Flox/Flox^Foxp3^YFP-Cre^* Treg cells (*SI Appendix* Fig. S5*C*) — its mRNA level was increased by 7828 folds (*SI Appendix* Fig. S5*D*). We confirmed the upregulation of CAI in *Cdc42^Flox/+^Foxp3^YFP-Cre^* Treg cells by quantitative real-time RT-PCR (Fig. 3*A*). CAI functions to modulate cellular pH by catalyzing the hydration of CO_2_ to HCO_3_^-^ and H^+^ (16). We predicted that the upregulation of CAI in *Cdc42^Flox/+^Foxp3^YFP-Cre^* Treg cells altered cellular pH, leading to Treg cell plasticity and anti-tumor T cell immunity. To test this, we measured the medium pH of *Cdc42^Flox/+^Foxp3^YFP-Cre^* and *Cdc42^+/+^Foxp3^YFP-Cre^* Treg cell culture. We found that while the pH of the *Cdc42^+/+^Foxp3^YFP-Cre^* Treg cell culture was maintained at 7.40, the pH of the *Cdc42^Flox/+^Foxp3^YFP-Cre^* Treg cell culture was changed to 7.60 (Fig. 3*B*). Importantly, *Cdc42^+/+^Foxp3^YFP-Cre^* Treg cells became plastic upon incubation with culture medium of pH 7.60 (Fig. 3 *C-E*) or with conditional medium from *Cdc42^Flox/+^Foxp3^YFP-Cre^* Treg cell culture (Fig. 3 *F-H*). The extracellular alkalization and plasticity of *Cdc42^Flox/+^Foxp3^YFP-Cre^* Treg cells were rescued by treatment with a CA inhibitor, acetazolamide (Fig. 3 *I-K*), and by CAI knockdown (Fig. 3 *L-O*). Thus, the plasticity of *Cdc42^Flox/+^Foxp3^YFP-Cre^* Treg cells is likely caused by CAI-mediated extracellular alkalization. Acetazolamide treatment of *Cdc42^Flox/+^Foxp3^YFP-Cre^* mice were able to revert tumor growth (Fig. 3*P*), correlating with reduced plasticity of intratumoral Treg cells and decreased tumorinfiltrating effector T cells *(SI Appendix* Fig. S6 *A-D*). We conclude that heterozygous *Cdc42* deletion induces Treg cell plasticity and revokes anti-tumor T cell immunity through upregulation of CAI. Of note, inhibition of CAI in *Cdc42^+/+^Foxp3^YFP-Cre^* Treg cells caused extracellular alkalization and Treg cell plasticity (Fig. 3 *I-O*), similar to that caused by the upregulation of CAI in *Cdc42^Flox/+^Foxp3^YFP-Cre^* Treg cells (Fig. 3 *A, I-K* and *M-O*). Furthermore, *Cdc42^+/+^Foxp3^YFP-Cre^* mice treated with acetazolamide showed similar tumor suppression to *Cdc42^Flox/+^Foxp3^YFP-Cre^* mice (Fig. 3*P*). These data suggest that physiological levels of CAI are essential for inhibition of Treg cell plasticity and for Treg cell-mediated tumor growth.

**Fig. 3.**
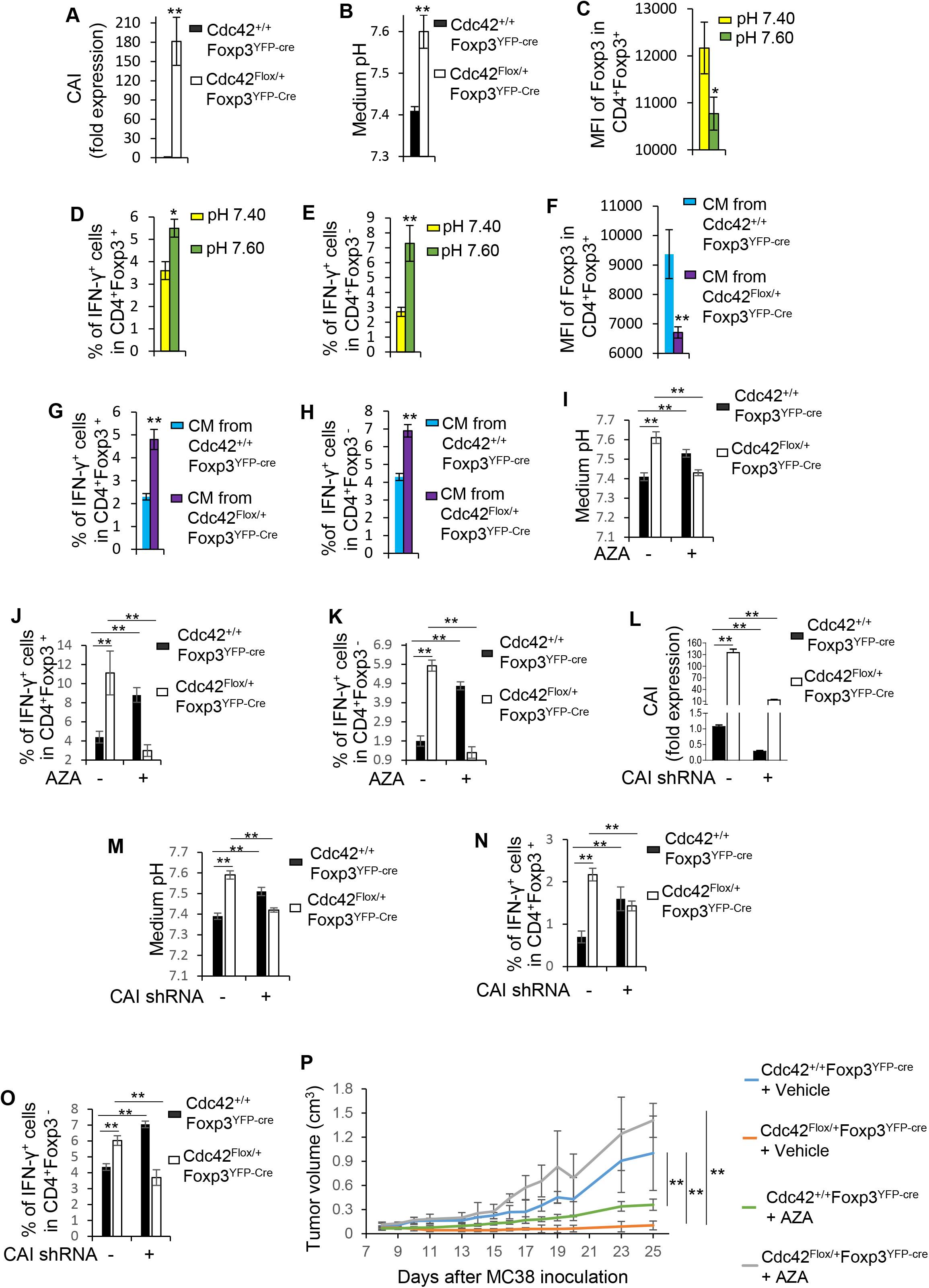
Heterozygous loss of *Cdc42* induces Treg cell plasticity and inhibits tumor growth through upregulation of CAI. (*A*) Quantitative real-time RT-PCR analysis of the expression of CAI mRNA in Treg cells from *Cdc42^+/+^Foxp3^YFP-Cre^* and *Cdc42^Flox/+^Foxp3^YFP-Cre^* mice. (*B*) Extracellular pH of *Cdc42^+/+^Foxp3^YFP-Cre^* and *Cdc42^Flox/+^Foxp3^YFP-Cre^* Treg cells cultured *ex vivo*. (*C* and *D*) Flow cytometry analysis of the expression of Foxp3 (*C*) and IFN-γ (*D*) in *Cdc42^+/+^Foxp3^YFP-Cre^* Treg cells cultured with normal medium (pH 7.40) or medium of pH 7.60. (*E*) Flow cytometry analysis of the expression of IFN-γ in CD4^+^ effector T cells (ex-Treg) that were converted from *Cdc42^+/+^Foxp3^YFP-Cre^* Treg cells cultured with normal medium or medium of pH 7.60. (*F* and *G*) Flow cytometry analysis of the expression of Foxp3 (*F*) and IFN-γ (*G*) in *Cdc42^+/+^Foxp3^YFP-Cre^* Treg cells incubated with conditional medium (CM) from *Cdc42^+/+^Foxp3^YFP-Cre^* or *Cdc42^Flox/+^Foxp3^YFP-Cre^* Treg cell culture. *(H)* Flow cytometry analysis of the expression of IFN-γ in CD4^+^ effector T cells (ex-Treg) that were converted from *Cdc42^+/+^Foxp3^YFP-Cre^* Treg cells incubated with conditional medium (CM) from *Cdc42^+/+^Foxp3^YFP-Cre^* or *Cdc42^Flox/+^Foxp3^YFP-Cre^* Treg cell culture. (*I*) Extracellular pH of *Cdc42^+/+^Foxp3^YFP-Cre^* and *Cdc42^Flox/+^Foxp3^YFP-Cre^* Treg cells cultured with or without AZA. (*J*) Flow cytometry analysis of the expression of IFN-γ in *Cdc42^+/+^Foxp3^YFP-Cre^* and *Cdc42^Flox/+^Foxp3^YFP-Cre^* Treg cells cultured with or without AZA. (*K*) Flow cytometry analysis of the expression of IFN-γ in CD4^+^ effector T cells (ex-Treg) that were converted from *Cdc42^+/+^Foxp3^YFP-Cre^* or *Cdc42^Flox/+^Foxp3^YFP-Cre^* Treg cells cultured with or without AZA. (*L*) Quantitative real-time RT-PCR analysis of the expression of CAI mRNA in *Cdc42^+/+^Foxp3^YFP-Cre^* and *Cdc42^Flox/+^Foxp3^YFP-Cre^* Treg cells transduced with or without CAI shRNA. (*M*) Extracellular pH of *Cdc42^+/+^Foxp3^YFP-Cre^* and *Cdc42^Flox/+^Foxp3^YFP-Cre^* Treg cells transduced with or without CAI shRNA. (*N*) Flow cytometry analysis of the expression of IFN-γ in *Cdc42^+/+^Foxp3^YFP-Cre^* and *Cdc42^Flox/+^Foxp3^YFP-Cre^* Treg cells transduced with or without CAI shRNA. (*O*) Flow cytometry analysis of the expression of IFN-γ in CD4^+^ effector T cells (ex-Treg) that were converted from *Cdc42^+/+^Foxp3^YFP-Cre^* or *Cdc42^Flox/+^Foxp3^YFP-Cre^* T reg cells transduced with or without CAI shRNA. (*P*) MC38 tumor growth in *Cdc42^+/+^Foxp3^YFP-Cre^* and *Cdc42^Flox/+^Foxp3^YFP-Cre^* mice treated with or without AZA. (*A*) Error bars indicate SD of 4 mice. (*B-O*) Error bars indicate SD of triplicates. Data are from 5 mice pooled. (*P*) Error bars indicate SD of 4 mice. Data are representative of two independent experiments. *p < 0.05; **p < 0.01. AZA: acetazolamide. MFI: Mean fluorescence intensity.

To define the mechanism underlying heterozygous *Cdc42* deletion-induced CAI upregulation in Treg cells, we analyzed transcription factor binding sites at the *CAI* locus. We found that the *CAI* locus contained the GATA3 binding motif WGATAA (Fig. 4*A*) that was reported in Treg cells (19). Interestingly, GATA3 was upregulated and its binding to the *CAI* locus was enhanced in *Cdc42^Flox/+^Foxp3^YFP-Cre^* Treg cells (Fig. 4 *B* and *C*). We attempted to genetically delete *GATA3* in *Cdc42^Flox/+^Foxp3^YFP-Cre^* Treg cells to determine whether loss of GATA3 could reverse CAI expression. However, we could not obtain *Cdc42^Flox/+^GATA3^Flox/Flox^Foxp3^YFP-Cre^* compound mice, possibly due to that heterozygous *Cdc42* deletion aggravates Treg cell-specific homozygous *GATA3* deletion-induced inflammatory disorders (20). Nonetheless, treatment of *Cdc42^Flox/+^Foxp3^YFP-Cre^* Treg cells with a GATA3 inhibitor, pyrrothiogatain (21), or GATA3 shRNA significantly suppressed the increased CAI expression (Fig. 4 *D* and *G*) and reduced *Cdc42^Flox/+^Foxp3^YFP-Cre^* Treg cell plasticity (Fig. 4 *E, F, H* and *I*). This suggests that the increased GATA3 expression and/or its binding to the *CAI* locus contribute to the increased CAI expression and plasticity of *Cdc42^Flox/+^Foxp3^YFP-Cre^* Treg cells. Importantly, pyrrothiogatain treatment of *Cdc42^Flox/+^Foxp3^YFP-Cre^* mice restored tumor growth (Fig. 4*J*), correlating with reduced plasticity of intratumoral Treg cells and decreased tumor-infiltrating effector T cells *(SI Appendix* Fig. S7 *A-D*), indicating that heterozygous *Cdc42* deletion induces Treg cell plasticity and revokes antitumor T cell immunity attributable to GATA3 upregulation. Of note, inhibition of GATA3 in *Cdc42^+/+^Foxp3^YFP-Cre^* Treg cells caused Treg cell plasticity (Fig. 4 *E, F, H*, and *I*), similar to that caused by the upregulation of GATA3 in *Cdc42^Flox/+^Foxp3^YFP-Cre^* Treg cells (Fig. 4 *B, E, F, H,* and *I*). And *Cdc42^+/+^Foxp3^YFP-Cre^* mice treated with pyrrothiogatain showed comparable tumor suppression to *Cdc42^Flox/+^Foxp3^YFP-Cre^* mice (Fig. 4*J*). The data suggest that similar to CAI, physiological levels of GATA3 are critical for inhibition of Treg cell plasticity and for Treg cell-mediated tumor growth.

**Fig. 4.**
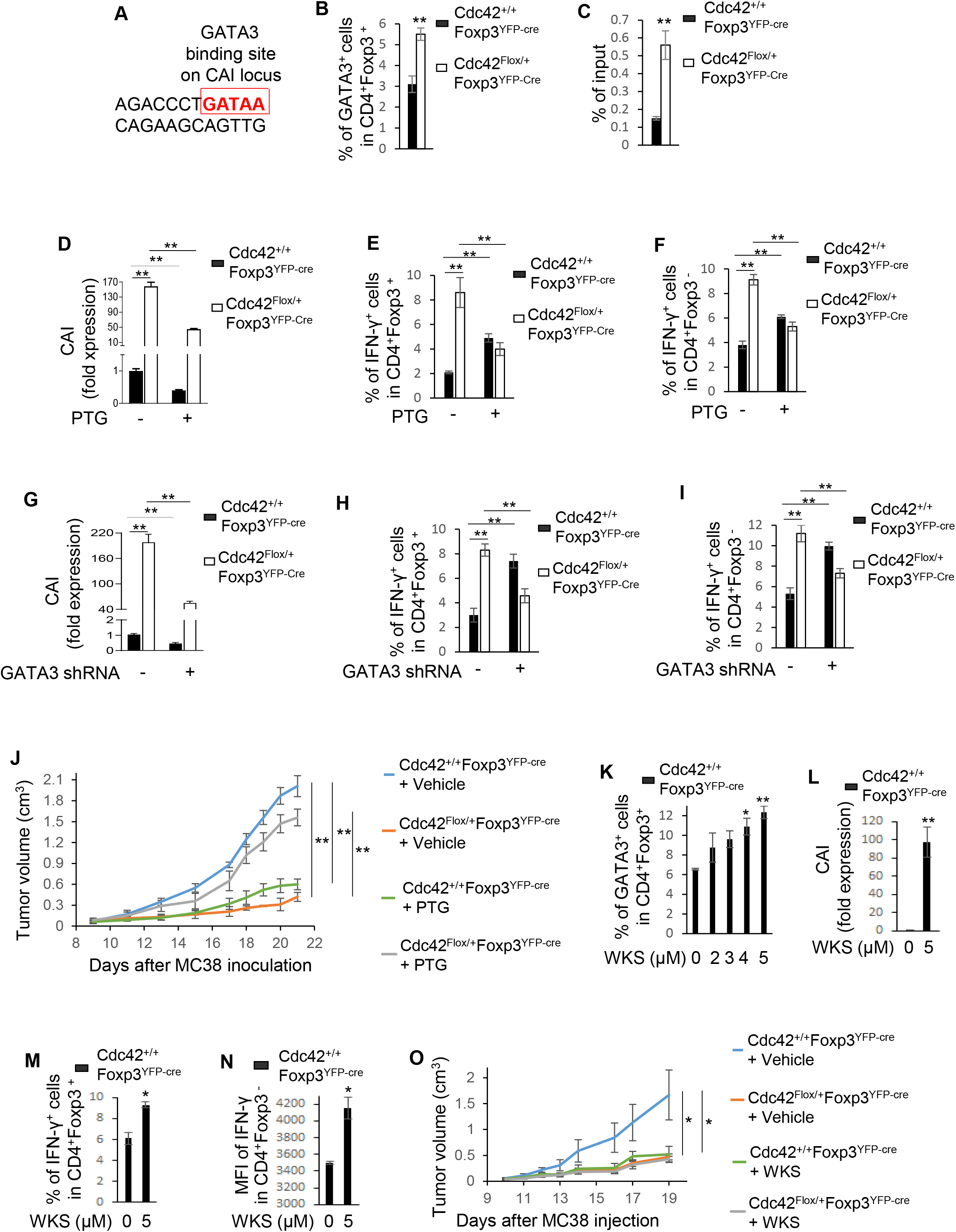
Heterozygous loss of *Cdc42* induces Treg cell plasticity and inhibits tumor growth through WASP-GATA3-mediated CAI expression. (*A*) Diagram of GATA3 binding sites on the CAI locus. (*B*) Flow cytometry analysis of the expression of GATA3 in Treg cells from *Cdc42^+/+^Foxp3^YFP-Cre^* and *Cdc42^Flox/+^Foxp3^YFP-Cre^* mice. (*C*) CHIP-qPCR analysis of GATA3 binding to the CAI locus in *Cdc42^+/+^Foxp3^YFP-Cre^* and *Cdc42^Flox/+^Foxp3^YFP-Cre^* Treg cells. (*D*) Quantitative real-time RT-PCR analysis of the expression of CAI mRNA in *Cdc42^+/+^Foxp3^YFP-Cre^* and *Cdc42^Flox/+^Foxp3^YFP-Cre^* Treg cells treated with or without PTG. (*E*) Flow cytometry analysis of the expression of IFN-γ in *Cdc42^+/+^Foxp3^YFP-Cre^* and *Cdc42^Flox+^Foxp3^YFP-Cre^* Treg cells treated with or without PTG. *(F)* Flow cytometry analysis of the expression of IFN-γ in CD4^+^ effector T cells (ex-Treg) that were converted from *Cdc42^+/+^Foxp3^YFP-Cre^* or *Cdc42^Flox/+^Foxp3^YFP-Cre^* Treg cells treated with or without PTG. (*G*) Quantitative real-time RT-PCR analysis of the expression of CAI mRNA in *Cdc42^+/+^Foxp3^YFP-Cre^* and *Cdc42^Flox/+^Foxp3^YFP-Cre^* Treg cells transduced with or without GATA3 shRNA. (*H*) Flow cytometry analysis of the expression of IFN-γ in *Cdc42^+/+^Foxp3^YFP-Cre^* and *Cdc42^Flox/+^Foxp3^YFP-Cre^* Treg cells transduced with or without GATA3 shRNA. (*I*) Flow cytometry analysis of the expression of IFN-γ in CD4^+^ effector T cells (ex-Treg) that were converted from *Cdc42^+/+^Foxp3^YFP-Cre^* or *Cdc42^Flox/+^Foxp3^YFP-Cre^* Treg cells transduced with or without GATA3 shRNA. (*J*) MC38 tumor growth in *Cdc42^+/+^Foxp3^YFP-Cre^* and *Cdc42^Flox/+^Foxp3^YFP-Cre^* mice treated with or without PTG. (*K*) Flow cytometry analysis of the expression of GATA3 in *Cdc42^+/+^Foxp3^YFP-Cre^* Treg cells treated with or without WKS. (*L*) Quantitative real-time RT-PCR analysis of the expression of CAI mRNA in *Cdc42^+/+^Foxp3^YFP-Cre^* Treg cells treated with or without WKS. (*M*) Flow cytometry analysis of the expression of IFN-γ in *Cdc42^+/+^Foxp3^YFP-Cre^* Treg cells treated with or without WKS. (*N*) Flow cytometry analysis of the expression of IFN-γ in CD4^+^ effector T cells (ex-Treg) that were converted from *Cdc42^+/+^Foxp3^YFP-Cre^* Treg cells treated with or without WKS. (*O*) MC38 tumor growth in *Cdc42^+/+^Foxp3^YFP-Cre^* and *Cdc42^Flox+^Foxp3^YFP-Cre^* mice treated with or without WKS. 1.4 x 10^6^ MC38 cells were injected. (*B*) Error bars indicate SD of 4 mice. (*C*) Error bars indicate SD of triplicates. Data are from 8 mice pooled. *(D-I* and *K-N)* Error bars indicate SD of triplicates. Data are from 5-6 mice pooled. (*J* and *O*) Error bars indicate SD of 4 mice. Data are representative of two independent experiments. *p < 0.05; **p < 0.01. PTG: pyrrothiogatain. WKS: wiskostatin. MFI: Mean fluorescence intensity.

We next examined how heterozygous *Cdc42* deletion upregulates GATA3 in Treg cells. Cdc42 functions through activating its immediate downstream effectors including WASP (22). Similar to heterozygous *Cdc42* deletion, genetic deletion of *WASP* has been shown to increase GATA3 expression in Treg cells (23). Consistently, we found that treatment of Treg cells with a WASP inhibitor, wiskostatin (24), enhanced the expression of GATA3 in a dose-dependent manner (Fig. 4*H*). Furthermore, wiskostatin treatment upregulated CAI (Fig. 4*I*) and increased Treg cell plasticity (Fig. 4 *J* and *K*). Consequently, inhibition of WASP by wiskostatin mimicked heterozygous *Cdc42* deletion in eliciting anti-tumor T cell immunity (Fig. 4*L* and *SI Appendix* Fig. S7 *E-H).* Collectively, these findings reveal that heterozygous *Cdc42* deletion induces Treg cell plasticity to promote anti-tumor T cell immunity through a WASP-GATA3-CAI signaling node.

### Pharmacological Targeting of Cdc42 Triggers Anti-tumor T Cell Immunity through Induction of Treg Cell Plasticity

To evaluate the potential of targeting Cdc42 signaling as a cancer immunotherapy, we examined whether pharmacologic inhibition of Cdc42 by a Cdc42 activity-specific inhibitor, CASIN (17), could mimic heterozygous *Cdc42* deletion in inducing Treg cell plasticity without causing systemic autoimmunity. As shown in Fig. 5, C57BL/6 mice treated with 30 mg/kg of CASIN twice a day for one week and then 40 mg/kg once a day for another week exhibited intact Treg cell homeostasis (Fig. 5*A*) but increased IFN-γ and/or IL-17 expression in Treg cells (Fig. 5*B*) and effector T cells (Fig. 5 *C* and *D*). Intriguingly, the mice did not display weight loss (Fig. 5*E*), tissue inflammation (Fig. 5*F*) and increased autoantibodies (Fig. 5*G*). Thus, at a certain treatment regimen, CASIN can recapitulate the phenotypes of heterozygous *Cdc42* deletion in Treg cells.

**Fig. 5.**
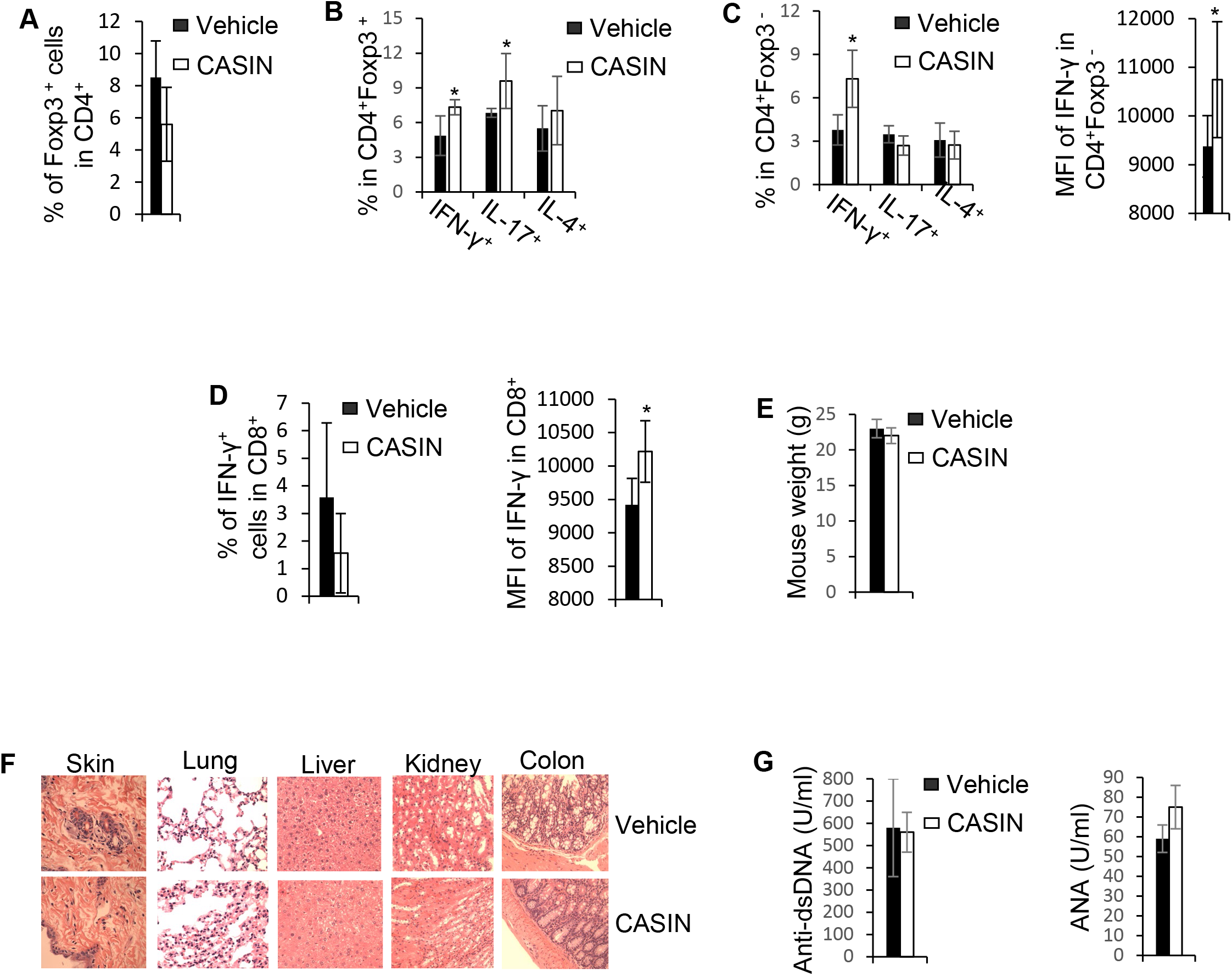
CASIN induces Treg cell plasticity without causing systemic autoimmune responses. (*A*) Flow cytometry analysis of the percentages of splenic Treg cells in C57BL/6 mice treated with or without CASIN. (*B-D*) Flow cytometry analysis of the expression of IFN-γ, IL-17 and/or IL-4 in Treg cells (*B*), CD4^+^ effector T cells (*C*) and CD8^+^ effector T cells (*D*). (*E*) Body weight of C57BL/6 mice. *(F)* H&E staining of the indicated organs. (*G*) ELISA analysis of the concentrations of serum autoantibodies. (*A-E*) and (*G*) Error bars indicate SD of 5 mice. *p < 0.05. (*F*) Data are representative of 3 mice. Anti-dsDNA: Anti-double stranded DNA. ANA: Antinuclear antibody. MFI: Mean fluorescence intensity.

Next, we prophylactically and therapeutically treated tumor-bearing mice with CASIN and found that CASIN effectively inhibited tumor growth (Fig. 6 *A* and *E*), correlating with intratumoral Treg cell platicity and increased effector T cells (Fig. 6 *B-D* and *F-H*). Depletion of T cells by anti-CD4/CD8 neutralizing antibodies completely restored tumor growth in CASIN-treated mice (Fig. 6*I*), suggesting that the tumor suppression in CASIN-treated mice is due to the increased effector T cell function. CASIN lost its inhibitory effect on tumor growth in *Cdc42^Flox/+^Foxp3^YFP-Cre^* mice (Fig. 6*J*). Together with our previous findings indicating that CASIN unlikely enhances effector T cell function in a cell-autonomous manner (25), the results implicate that CASIN triggers antitumor T cell immunity through induction of Treg cell plasticity. To substantiate this, we directly induced Treg cell plasticity *ex vivo* with CASIN (Fig. 7 *A* and *B*) at concentrations equivalent to the serum doses in therapeutic CASIN-treated tumor-bearing mice (*SI Appendix* Fig. S8). Upon prophylactic (Fig. 7*C*) and therapeutic (Fig. 7*I*) transfer with congenic (CD45.1^+^) effector T cells into *Rag1^-/-^* mice, these CASIN-treated CD45.2^+^ Treg cells diminished tumor growth (Fig. 7 *D* and *J*). The tumor suppression was associated with a substantial increase in tumor-infiltrating Treg cell plasticity (Fig. 7 *E* and *K*) and effector T cells that were converted from (Fig. 7 *F* and *L*) or unleashed by (Fig. 7 *G, H, M* and *N*) plastic Treg cells. CASIN did not cause systemic autoimmunity in tumor-bearing mice (*SI Appendix* Fig. S9) similar to that in tumor-free mice, supporting that CASIN administration does not elicit off-target and off-Treg cell effects in suppressing tumor growth and allows for a therapeutic window for cancer treatment.

**Fig. 6.**
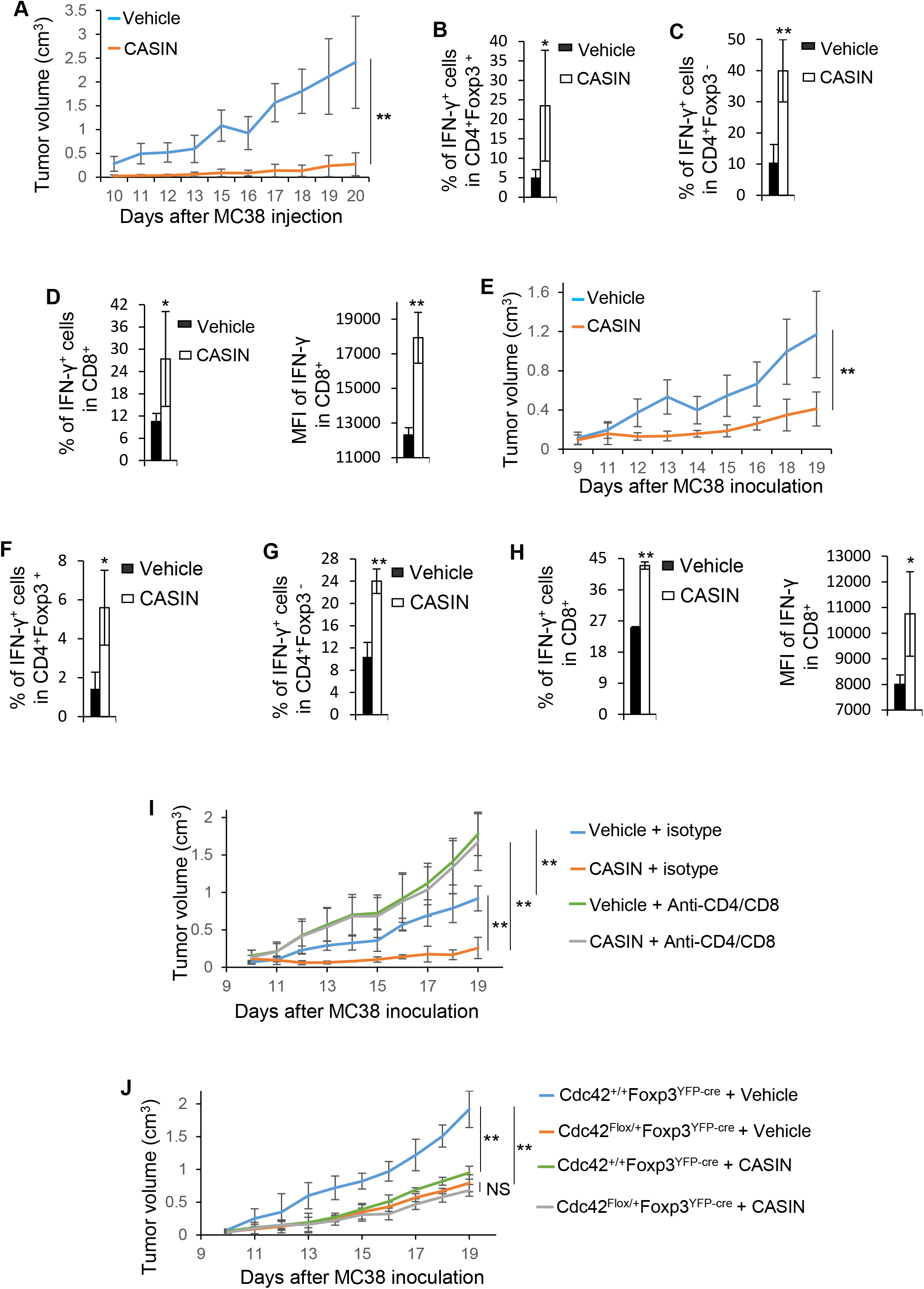
CASIN administration elicits anti-tumor T cell immunity. (*A*) MC38 tumor growth in C57BL/6 mice treated with or without CASIN starting at the same time as MC38 cell inoculation. (*B-D*) Flow cytometry analysis of the expression of IFN-γ in tumor-infiltrating Treg cells (*B*), CD4^+^ effector T cells (*C*) and CD8^+^ effector T cells (*D*) from C57BL/6 mice treated with or without CASIN starting at the same time as MC38 cell inoculation. (*E*) MC38 tumor growth in C57BL/6 mice treated with or without CASIN upon MC38 tumor onset. *(F-H)* Flow cytometry analysis of the expression of IFN-γ in tumor-infiltrating Treg cells (*F*), CD4^+^ effector T cells (*G*) and CD8^+^ effector T cells (*H*) from C57BL/6 mice treated with or without CASIN starting upon MC38 tumor onset. (*I*) MC38 tumor growth in C57BL/6 mice treated with or without Anti-CD4/CD8 neutralizing antibodies and/or CASIN starting upon MC38 tumor onset. (*J*) MC38 tumor growth in *Cdc42^+/+^Foxp3^YFP-Cre^* and *Cdc42^Flox/+^Foxp3^YFP-Cre^* mice treated with or without CASIN starting upon MC38 tumor onset. 1.4 x 10^6^ MC38 cells were injected. Error bars indicate SD. n = 5 (*I* and *J)* or 6 *(A-H).* Data are representative of two independent experiments. *p < 0.05; **p < 0.01. MFI: Mean fluorescence intensity. NS: No significance.

**Fig. 7.**
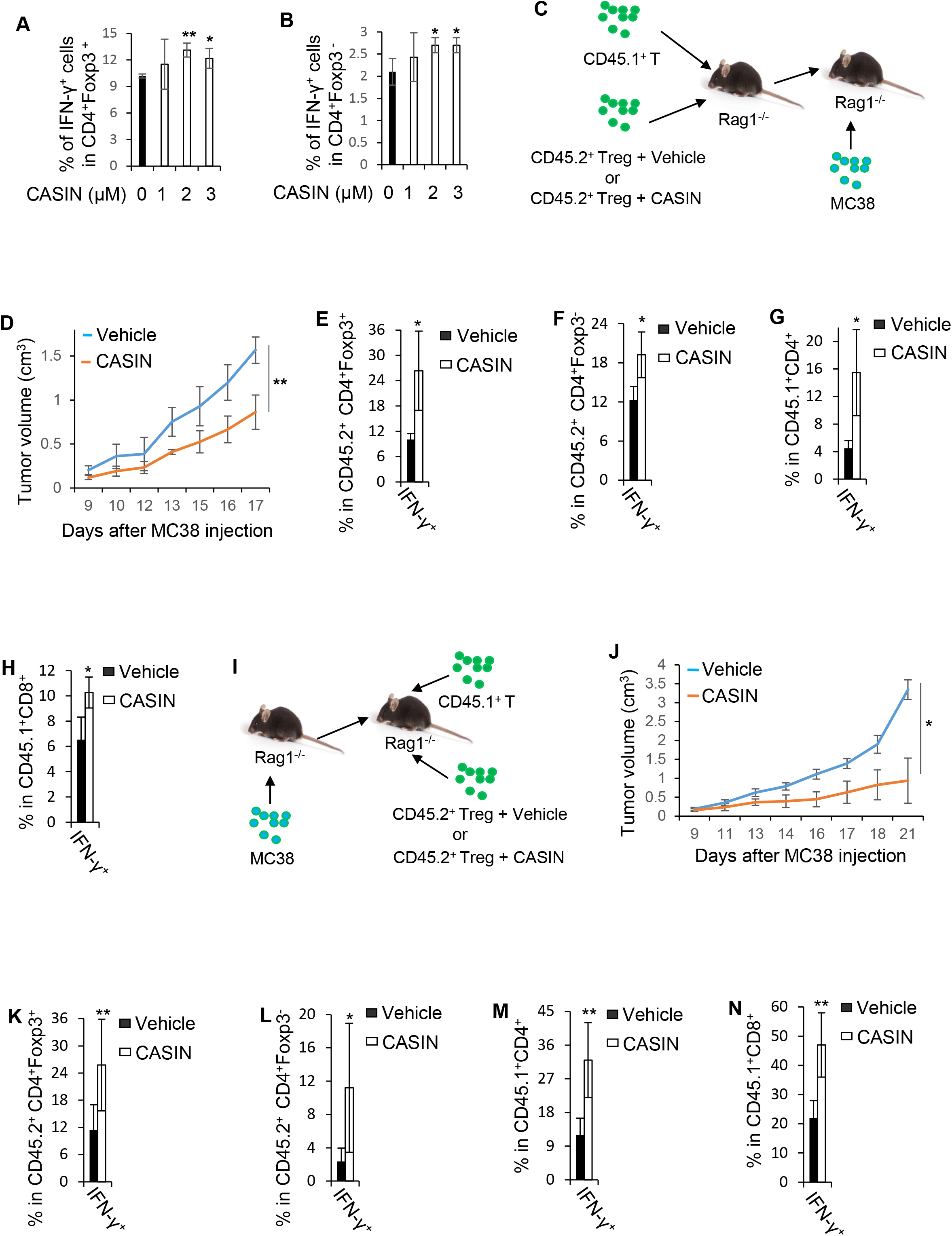
CASIN elicits anti-tumor T cell immunity through induction of Treg cell plasticity. (*A* and *B*) Flow cytometry analysis of the expression of IFN-γ in Treg cells (*A*) and CD4^+^ effector T cells (ex-Treg) (*B*) that were converted from Treg cells treated with or without CASIN *ex vivo.* (*C*) Schematic overview of prophylactic co-transfer of Treg cells treated with or without CASIN (2 μM) and congenically labelled effector T cells. (*D*) MC38 tumor growth in the recipient mice prophylactically co-transferred with Treg cells treated with or without CASIN and effector T cells. *(E-H)* Flow cytometry analysis of the expression of IFN-γ in tumor-infiltrating Treg (CD45.2^+^CD4^+^Foxp3^+^) (*E*), ex-Treg (CD45.2^+^CD4^+^Foxp3^-^) (*F*), and congenically labelled effector T cells (CD45.1^+^CD4^+^, CD45.1^+^CD8^+^) (*G* and *H*) in the recipient mice prophylactically co-transferred with Treg cells treated with or without CASIN and effector T cells. (*I*) Schematic overview of therapeutic co-transfer of Treg cells treated with or without CASIN (2 μM) and congenically labelled effector T cells. (*J*) MC38 tumor growth in the recipient mice therapeutically co-transferred with Treg cells treated with or without CASIN and effector T cells. *(K-N)* Flow cytometry analysis of the expression of IFN-γ in tumor-infiltrating Treg (CD45.2^+^CD4^+^Foxp3^+^) (*K*), ex-Treg (CD45.2^+^CD4^+^Foxp3^-^) (*L*), and congenically labelled effector T cells (CD45.1^+^CD4^+^, CD45.1^+^CD8^+^) (*M* and *N)* in the recipient mice therapeutically co-transferred with Treg cells treated with or without CASIN and effector T cells. (*A* and *B*) Data are from 5 mice pooled. Error bars indicate SD of triplicates. (*D-H*) Error bars indicate SD of 4 mice. (*J-N*) Error bars indicate SD of 7 mice. *p < 0.05; **p < 0.01.

### Pharmacological Targeting of Cdc42 Potentiates ICI Activity in Eliciting Anti-tumor T Cell Immunity

Although ICIs have achieved clinical success in several types of cancer, primary or acquired resistance in the majority of cancer patients limited their broad application (7). We thus examined whether CASIN can potentiate ICIs in tumor suppression. We found that combined CASIN and Anti-PD-1 treatment inhibited MC38 tumor growth to a greater extent than single treatment (Fig. 8*A*). Combined CASIN and Anti-PD-1 treatment did not further increase Treg cell plasticity, compared to CASIN alone (Fig. 8*B*). However, the combinatory treatment led to more effector T cells (Fig. 8 *C* and *D*), likely due to the ability of Anti-PD-1 to reinvigorate exhausted effector T cells. Moreover, compared to single treatment, combined CASIN and Anti-PD-1 led to greater suppression of tumor growth of human colon cancer cells, HCT116, in mouse xenografts engrafted with human CD34^+^ hematopoietic stem cells (Fig. 8*E*). CASIN induced human Treg cell plasticity. Such plasticity was not further increased by the combinatory treatment (Fig. 8*F*). However, the combinatory treatment further increased human effector T cells (Fig. 8 *G* and *H*). These results suggest that CASIN potentiates ICIs in eliciting anti-tumor T cell immunity.

**Fig. 8.**
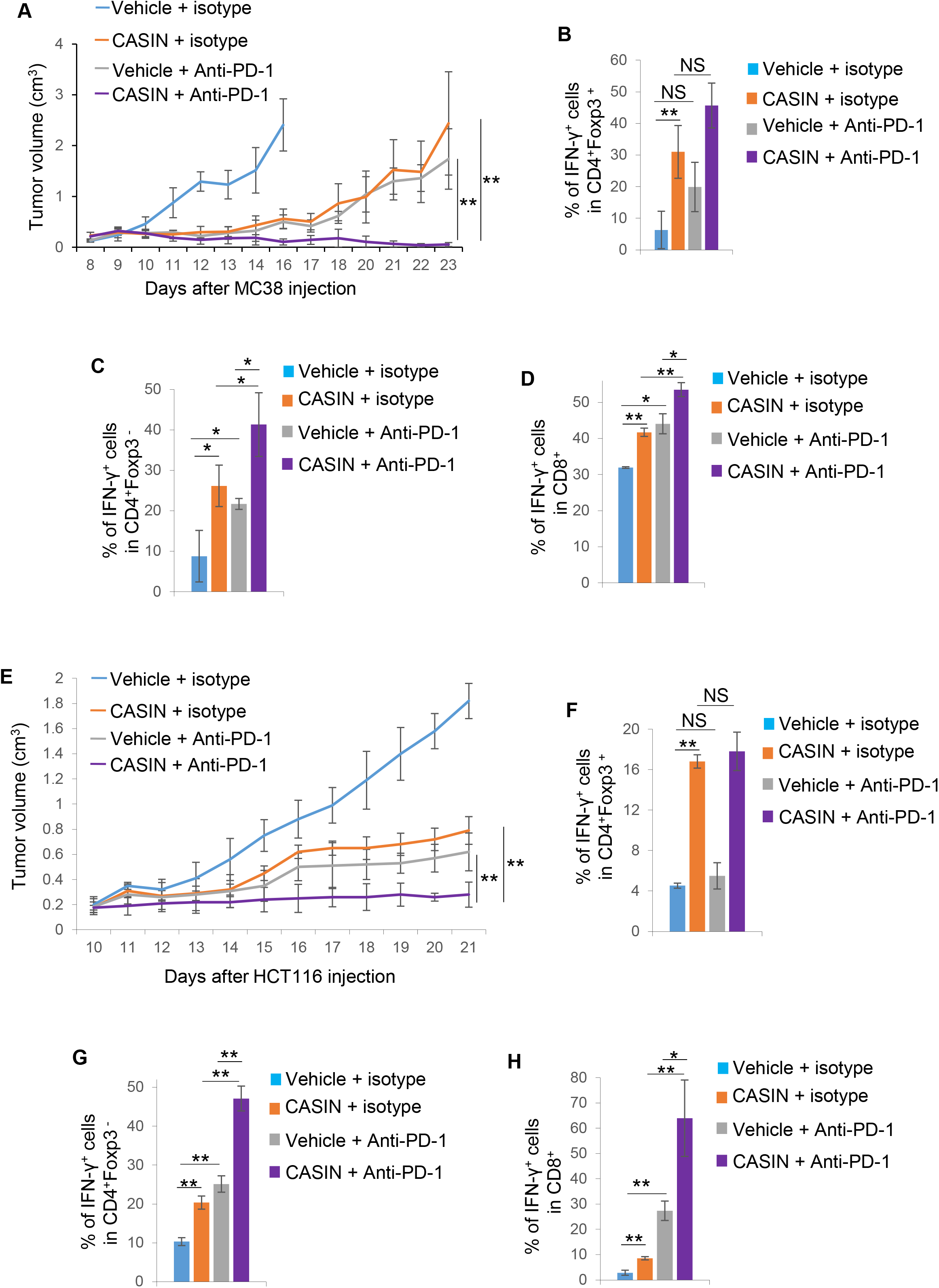
CASIN enhances anti-PD-1-induced anti-tumor T cell immunity. (*A*) MC38 tumor growth in C57BL/6 mice treated with or without Anti-PD-1 and/or CASIN starting upon MC38 tumor onset. (*B-D*) Flow cytometry analysis of the expression of IFN-γ in MC38 tumor-infiltrating Treg (*B*) and effector T cells (*C* and *D*). (*E*) HCT116 tumor growth in NSGS mice treated with or without Anti-PD-1 and/or CASIN starting on HCT116 tumor onset. *(F-H)* Flow cytometry analysis of the expression of IFN-γ in HCT116 tumor-infiltrating Treg (*F*) and effector T cells (*G* and *H*). Error bars indicate SD. n = 3 (*A-D*) or 4 (*E-H*). Data are representative of two independent experiments. *p < 0.05; **p < 0.01. NS: No significance.

### Pharmacological Targeting of Cdc42 Aggravates Colitis

CASIN treatment of steady-state and tumor-bearing mice did not cause spontaneous inflammatory disorders (Fig. 5*E-G* and *SI Appendix* Fig. S9). However, CASIN treatment of mice bearing colitis that was induced by dextran sulfate sodium (DSS) led to more weight loss, compared to vehicle treatment (*SI Appendix* Fig. S10*A*). Consistent with an essential role of Th1 cells in the development of colitis (15), CASIN treatment increased the production of IFN-γ in colonic CD4^+^ effector T cells (*SI Appendix* Fig. S10*B*). CASIN induced Treg cell plasticity in the colon, as evidenced by elevated IFN-γ^+^ Treg cells (*SI Appendix* Fig. S10*C*). Given our previous finding that CASIN did not enhance Th1 cell differentiation (25), it is logical to reason that the increased IFN-γ^+^ effector T cells are attributable to Treg cell plasticity in CASIN-treated mice. These results caution the use of CASIN in cancer patients with autoimmune diseases.

## Discussion

In this study, we show that heterozygous deletion of *Cdc42* induces Treg cell plasticity, leading to anti-tumor T cell immunity, suggesting that Cdc42 is required for maintenance of Treg cell fitness and tumor immune evasion. Mechanistically, Cdc42 regulates Treg cell fitness and tumor immune evasion through a non-canonical signaling cascade, i.e. WASP-GATA3-CAI. Inhibition of WASP induces Treg cell plasticity and anti-tumor T cell immunity, mimicking heterozygous deletion of *Cdc42*. We provide data to indicate that WASP suppresses Treg cell plasticity and anti-tumor T cell immunity through restraining GATA3 expression. Given that WASP can translocate to the nucleus to regulate gene transcription in Th1 cells (26), it is plausible that the nuclear function of WASP is involved in regulating GATA3 expression in Treg cells. We show that GATA3 expression in Treg cells contributes to heterozygous Cdc42 deletion-induced Treg cell plasticity (e.g. effector T cell cytokine IFN-γ expression in Treg cells) and anti-tumor T cell immunity. As GATA3 is known to inhibit IFN-γ expression in effector T cells (e.g. Th2 cells), our observation that GATA3 promotes IFN-γ expression in immunosuppressive Treg cells suggests that GATA3 modulates IFN-γFexpression in a cell type-dependent manner. We implicate that GATA3 may promote Treg cell plasticity/IFN-γ expression in Treg cells through upregulation of CAI. CAI is expressed in the cytoplasm, and the increased CAI in heterozygous *Cdc42* knockout Treg cells is expected to elevate intracellular HCO_3_^-^ and H^+^. The alkalization of the extracellular pH in heterozygous *Cdc42* knockout Treg cell culture suggests that heterozygous *Cdc42* deletion facilitates HCO_3_^-^ transportation into the extracellular space, which could result in intracellular accumulation of H^+^ and thus acidification of heterozygous *Cdc42* knockout Treg cells. Such alkalization of extracellular pH can induce Treg cell plasticity, enabling effector T cells to fight against tumor growth. How does alkalization of extracellular pH destabilize Cdc42^+/-^ Treg cells remains elusive. We speculate that alkali-sensing receptors (e.g. InsR-RR) sense extracellular alkaline (27) and then elicit intracellular signals to promote DNMT3a expression and thus methylation of CNS2 of the Foxp3 locus, leading to loss of stable Foxp3 expression and subsequent derepression of effector T cell cytokine (e.g. IFN-γ) expression.

Pharmacological targeting of Cdc42 by CASIN recapitulates heterozygous *Cdc42* deletion-induced phenotypes in Treg cell plasticity and anti-tumor T cell responses without causing autoimmunity. Interestingly, Treg cell plasticity induced by transient CASIN treatment *in vitro* triggers anti-tumor T cell immunity *in vivo,* suggesting that an *ex vivo* CASIN treatment strategy could be useful for adoptive Treg/T cell transfer to treat cancer, avoiding potential systemic toxicity from *in vivo* administration of the inhibitor.

We have reported that homozygous *Cdc42* deletion in Treg cells decreases Treg cell numbers, induces Treg cell plasticity, and causes systemic inflammatory disorders (15). In this study, we show that heterozygous *Cdc42* deletion also induces Treg cell plasticity but does not alter Treg cell homeostasis and cause spontaneous autoimmunity. Thus, the autoimmune responses in homozygous *Cdc42* knockout mice are presumably caused by dampened Treg cell homeostasis but not Treg cell plasticity. Of note, although heterozygous *Cdc42* deletion appears to downregulate Foxp3 expression to an extent comparable to homozygous *Cdc42* deletion, it causes milder effector T cell reprogramming in Treg cells. Interestingly, CASIN treatment mimics heterozygous but not homozygous *Cdc42* deletion in modulating Treg cells. Importantly, CASIN treatment also does not trigger autoimmune diseases. It is noteworthy mentioning that while Treg cell plasticity induced by heterozygous *Cdc42* deletion and CASIN is moderate in steady-state mice, the plasticity phenotypes are magnified in the tumor microenvironment in heterozygous *Cdc42* knockout and CASIN-treated tumor-bearing mice. We provide data to indicate that not only the magnified Treg cell plasticity but also dampened Treg cell suppressive function contribute to tumor suppression. Similar to that in steady-state mice, CASIN treatment does not cause autoimmunity in tumor-bearing mice. These findings collectively suggest that the dosage effect of Cdc42 expression/activity in Treg cells can be harnessed to unleash anti-tumor T cell immunity without causing autoimmunity.

It is debatable why heterozygous *Cdc42* deletion causes gene dosage effect and why CASIN mimics heterozygous but not homozygous *Cdc42* deletion. To begin to understand this, we have examined the activity of WASP and PAK1, two immediate downstream effectors of Cdc42, in heterozygous and homozygous *Cdc42* knockout and CASIN-treated Treg cells. Interestingly, heterozygous *Cdc42* deletion causes milder inhibition of WASP and PAK1 activation and CASIN exhibits inhibitory effect on WASP and PAK1 activation to a similar extent to heterozygous but not homozygous *Cdc42* deletion (*SI Appendix* Fig. S11). Hence, the dosage effect of heterozygous *Cdc42* deletion and CASIN seems to stem from their dosage effect on WASP and PAK1.

Cdc42 is known to promote tumorigenesis (28). Overexpression, hyper-activation, and/or recurrent mutations of Cdc42 have been associated with poor prognosis in multiple human cancers (29, 30). Thus, CASIN may have tumor cell-intrinsic effect in suppressing tumor growth. However, the complete restoration of tumor growth by T cell depletion in CASIN-treated mice indicates that the immuno-effect outweighs any tumor cell-intrinsic effect, suggesting that Cdc42 may serve as an immunotherapeutic target. Nevertheless, given multiple signaling roles of Cdc42 in cell biology, whether Cdc42 targeting will allow for a therapeutic window comparing immunomodulation benefit versus toxicity needs a careful examination. In this aspect, although overexpression of constitutively active and dominant negative Cdc42 mutants showed broad functions of Cdc42 in a variety of cell types, a more specific and physiological approach of genetic deletion of Cdc42 has revealed limited effects of Cdc42 suppression in primary cells including hematopoietic stem and progenitor cells, fibroblastoid cells, and B cells (31–33). For example, contrary to the prevailing view that Cdc42 promotes cell survival, gene deletion of Cdc42 did not affect survival of hematopoietic stem cells. Thus, effective and specific Cdc42 inhibitors that dose-dependently suppress Cdc42 activity are expected to have manageable toxicity. In support of this, CASIN appears to specifically and reversibly inhibit Cdc42 activity at a dose range tested (17) without detectable effect on thymocyte development, T cell homeostasis, and Th1/Th17 cell differentiation *in vitro* and *in vivo* (25). CASIN does not appear to have off-target and off-Treg cell effects nor does it cause systemic inflammation in either tumor-free or tumor-bearing mice. Therefore, a regimen of Cdc42 inhibitor administration, with an optimal profile of toxicity, pharmacokinetics and pharmacodynamics, could present a therapeutic window for Treg cell plasticity induction and subsequent anti-tumor T cell immunity. Of note, the Kd of CASIN for binding to Cdc42 is ~ 0.3 uM and the solubility is ~100 uM (17), leaving room for further optimization.

## Materials and Methods

### Mice

*Foxp3^YFP-Cre^* (016959, RRID:IMSR_JAX:016959), *Foxp3*^eGFP-Cre-ERT2^ (016961, RRID:IMSR_JAX:016961), *Rosa26^eYFP^* (006148, RRID:IMSR_JAX:006148), C57BL/6/J (000664, RRID:IMSR_JAX:000664), and *Rag1*^-/-^ (002216, RRID:IMSR_JAX:002216) mice were purchased from Jackson laboratories. *Cdc42^Flox/Flox^* mice were generated as described previously (34). *Cdc42^Flox/Flox^* mice were bred with *Foxp3^YFP-Cre^* mice in our animal facility to generate *Cdc42^Flox/+^Foxp3^YFP-Cre^* mice. *Foxp3*^eGFP-Cre-ERT2^ mice were bred with *Rosa26^eYFP^* mice to generate *Foxp3*^eGFP-Cre-ERT2^*Rosa26^eYFP^* mice. *Foxp3*^eGFP-Cre-ERT2^*Rosa26^eYFP^* mice were then bred with *Cdc42^Flox/Flox^* mice to generate *Cdc42^Flox/+^Foxp3*^eGFP-Cre-ERT2^*Rosa26^eYFP^* mice. NSGS mice were obtained from the Comprehensive Mouse and Cancer Core at the Cincinnati Children’s Hospital Medical Center. NSGS mice engrafted with human CD34^+^ hematopoietic stem cells were generated as described previously (35). Unless otherwise noted, 6 to 8 weeks old male and female mice were used.

All mice were housed under specific pathogen-free conditions in the animal facility at the Cincinnati Children’s Hospital Research Foundation in compliance with the Cincinnati Children’s Hospital Medical Center Animal Care and Use Committee protocols.

### Tumor Cell Lines

Mouse colon adenocarcinoma GFP-tagged tumor cell line MC38 and human colon adenocarcinoma tumor cell line HCT116 were generously provided by Dr. Joseph Palumbo at the Cincinnati Children’s Hospital Medical Center. Mouse pancreatic cancer cell line KPC was generously provided by Dr. Matthew Flick at the Cincinnati Children’s Hospital Medical Center. All tumor cell lines were grown in DMEM with 10% fetal calf serum (FCS). Regular mycoplasma testing was carried out on all cells maintained in culture.

### Tumor Growth Studies and Treatments

Unless otherwise noted, 8 x 10^5^ MC38 or KPC tumor cells were injected subcutaneously in 100 μl PBS into one of the flanks of the indicated mice. Two million HCT116 cells were injected subcutaneously in 100 μl PBS into one flank of NSGS mice 15 weeks after the mice were transplanted with CD34^+^ hematopoietic stem cells. Wherever possible, animals were randomized into treatment groups. Tumor volumes were measured every day after tumor became visible and calculated as V = (length × width^2^) x 0.50. All the mice were euthanized when tumor volume of control mice reached about 2 cm^3^ and tumors were harvested.

CASIN (Cayman Chemical, 17694) in 60% PBS and 40% β-cyclodextrin (Cayman Chemical, 23387) was injected intraperitoneally twice daily at 30 mg/kg for 7 days and then once daily at 40 mg/kg until the end of the experiments. For prophylactic treatment, CASIN injection was started at the same time as tumor cell injection. For therapeutic treatment, CASIN injection was started at around day 10 when tumors became visible. CA inhibitor acetazolamide (Sigma-Aldrich, A6011) in 90% PBS, 5% Tween-20 and 5% polyethylene glycol was injected intraperitoneally once daily at 40 mg/kg starting at the same time as tumor cell injection. WASP inhibitor wiskostatin (Cayman Chemical, 15047) in 97.5% PBS and 2.5% DMSO was injected intraperitoneally once daily at 60 mg/kg starting at the same time as tumor cell injection. GATA3 inhibitor pyrrothiogatain (Santa Cruz, sc-352288A) in 90% PBS, 5% Tween-20 and 5% polyethylene glycol was injected intraperitoneally once daily at 60 mg/kg starting at the same time as tumor cell injection. For T cell depletion experiments, anti-CD4 (150 μg/mouse) (Bio X Cell, BE0003-3, Clone ID: YTS 177, RRID:AB_1107642) and anti-CD8 (150 μg/mouse) (Bio X Cell, BE0118, Clone ID: HB-129, RRID: AB_10949065) or rat IgG2a isotype control (150 μg/mouse) (Bio X Cell, BE0090, Clone ID: LTF-2, RRID:AB_1107780) were injected intraperitoneally every other day for 4 injections and then twice a week until the end of the experiments. For experiments involving ICIs, anti-PD1 (150 μg/mouse) (Bio X Cell, BE0146, Clone ID: RMP1-14, RRID: AB_10949053) was injected every other day for a total of 5 injections.

### Adoptive Cell Transfer Studies

Splenic Treg cells were obtained by magnetic-activated cell sorting of CD4^+^ T cells (Miltenyi, 130-117-043) followed by FACS sorting of CD4^+^YFP^+^ Treg cells to >99% purity. Splenic T cells were isolated from congenic BoyJ mice by a pan T cell isolation kit (Miltenyi, 130-095-130) followed by Treg cell depletion with CD25 MicroBead Kit (Miltenyi, 130-091-072). The sorted Treg cells (2-3 x 10^5^ per mouse) with or without congenic T cells (6 x 10^5^ per mouse) were transferred intraperitoneally into *Rag1^-/-^* mice 1 day before or ~ 10 days after MC38 cell injection into the recipient mice.

### Preparation of Single-cell Suspensions, Antibody Staining and Flow Cytometry

Spleens were mashed with a syringe plunger and passed through 40 μm cell strainer, followed by treatment with red blood cell lysis buffer (BD Biosciences, 555899). Tumors were minced into small fragments and treated with 1.5 mg/ml collagenase IV (Sigma, C5138) for 30 min at 37 °C under agitation. The digested tumor tissue was then filtered through a 70 μm cell strainer and centrifuged at 1500 rpm at 4 °C for 5 min. The pellets were dissolved in 8 ml of 40% percoll and slowly layered over 5 ml of 80% percoll in a 15 ml falcon tube. The falcon tube was then centrifuged at 2000 rpm at 4 °C for 20 min, stopping without brakes. Cells at the interface between 40% percoll and 80% percoll were carefully removed and washed twice with complete RPMI T-cell medium. The cells from spleens and tumors were re-stimulated with PMA and Ionomycin for 5 hrs in the presence of Golgi plug for the last 4 hrs, and then subjected to antibody staining. Cell surface proteins were stained for 20 min at 4 °C with the following antibodies: CD45.1 (BioLegend, 110714, Clone ID: A20, RRID:AB_313503), CD45.2 (BioLegend, 109825, Clone ID: 104, RRID:AB_893351), ICOS (eBioscience, 12-9942-81, Clone ID: 7E.17G9, RRID:AB_466273), PD-1 (eBioscience, 12-9985-81, Clone ID: J43, RRID:AB_466294), GITR (eBioscience, 25-5874-80, Clone ID: DTA-1, RRID:AB_10544396), CTLA-4 ( eBioscience, 12-1522-82, Clone ID: UC10-4B9, RRID:AB_465879), CD39 (eBioscience, 25-0391-82, Clone ID: 24DMS-1, RRID:AB_1210766), CD73 (eBioscience, 25-0731-80, Clone ID: TY/11.8, RRID:AB_10870789), CD4 (eBioscience, 48-0042-82, Clone ID: RM4-5, RRID:AB_1272194), and CD8α (eBioscience, 25-0081-82, Clone ID: 53-6.7, RRID:AB_469584). Intracellular proteins were stained for 60 min at room temperature after permeabilization and fixation with BD Cytofix/Cytoperm Plus (BD Biosciences, 555028) using the following antibodies: IL-4 (eBioscience, 12-7041-82, Clone ID: 11B11, RRID:AB_466156), IL-17A (BD Pharmingen, 559502, Clone ID: TC11-18H10, RRID:AB_397256), IFN-γ (BioLegend, 505810, 505826, Clone ID: XMG1.2, RRID:AB_315404, RRID:AB_2295770), Foxp3 (eBioscience, 17-5773-82, Clone ID FJK-16s, RRID:AB_469457), and GATA-3 (eBioscience, 25-9966-42, Clone ID: TWAJ, RRID:AB_2573568). The stained cells were analyzed by BD LSRII, FACSCanto, or LSRFortessa flow cytometers. Data were analyzed with BD FACSDiva.

In certain experiments, Treg cells isolated by FACS sorting of CD4^+^YFP^+^ cells to >99% purity were expanded with Treg cell expansion kit (Miltenyi, 130-095-925) for 3 days, in the presence or absence of 47.6 mM NaHCO_3_ (to make culture medium of pH 7.60), 3 μM acetazolamide, 50 μM pyrrothiogatain, or the indicated concentrations of wiskostatin or CASIN. The cells were then subjected to the indicated analysis. For cytokine detection, the cells were re-stimulated with PMA and ionomycin along with Golgi Plug followed by flow cytometry analysis. For lentiviral shRNA-mediated knockdown, scramble and CAI shRNA lentiviral supernatant were produced by transfection of 293T cells with packaging lentiviral plasmids and either scramble or CAI shRNA lentiviral vectors (Origene, TL510087) followed by concentrating with Lenti-X concentrator (Takara, 631231). The viral supernatants were transduced into Treg cells that were expanded for 1 day before the transduction, by centrifugation at 1000g at 32°C for 90 min. The transduced Treg cells were further expanded for another 2 days and subjected to the indicated analysis.

For 5-bromo-29-deoxyuridine (BrdU) incorporation assay, mice were injected intraperitoneally with 500 mg BrdU. Two hours after injection, splenocytes were isolated and immunolabeled with anti-CD4 and anti-Foxp3 antibodies and BrdU incorporation was analyzed by a BrdU Flow kit per the manufacturer’s protocol (BD Pharmingen, 552598) (15).

For cell apoptosis assay, freshly isolated splenocytes were immunolabeled with anti-CD4 and anti-Foxp3 antibodies and cell apoptosis was analyzed by anti-active caspase 3 antibody (BD Pharmingen, 559341, Clone ID: C92-605, RRID:AB_397234) staining followed by flow cytometry.

### Autoantibody Detection

Serum was isolated from blood using serum separator tubes. Anti-dsDNA and ANA in serum were detected by ELISA using the corresponding ELISA Kits (Alpha Diagnostics, 51d53s and 5212, respectively), according to the manufacturer’s protocol.

### Bisulfite Pyrosequencing of Methylation of Foxp3 Enhancer

A total of 200 ng genomic DNA was subjected to sodium bisulfite treatment and purified using the EZ DNA Methylation-Gold Kit (Zymo research, D5007) according to the manufacturer’s specifications. Two rounds of standard PCR amplification reaction were performed to amplify targeted gene fragment at an annealing temperature of 50° C before being subjected to pyro-sequencing. The generated pyrograms were automatically analyzed using PyroMark analysis software (Qiagen). The pyrosequencing assay was validated using SssI-treated human genomic DNA as a 100% methylation control and human genomic DNA amplified by GenomePlex Complete Whole Genome Amplification kit (Sigma-Aldrich, WGA2-50RXN) as 0% methylation control. Primers used for bisulfite pyrosequencing are as follows:

mFoxp3_assay1_NF: 5’-TTTGTGTTTTTGAGATTTTAAAATT-3’; mFoxp3_assay1_NR: /5Biosg/5’-AAAAATAA CTAATCTATCCTATAACC-3’; mFoxp3_assay1_LF: 5’-TATTTTTTTGGGTTTTGGGATATTA-3’; mFoxp3_ assay1_LR: 5’-ACAAATAATCTACCCCACAAATTTC-3’; mFoxp3_assay1_S2: 5’-GGGTTTTTTTGGTATTTAA GAAAG-3’; mFoxp3_assay1_S3: 5’-GGGTTTTGTATGGTAGTTAGATGG-3’; mFoxp3_assay1_S4: 5’-AGTATT TATATTATTTTATTTGGG-3’; mFoxp3_assay2_NF: 5’-GGTTATAGGATAGATTAGTTATTTTT-3’; mFoxp3_ assay2_NR: /5Biosg/5’-CCAACTTCCTACACTATCTATTAAAAC-3’; mFoxp3_assay2_LF: 5’-TTTATATTATTTT ATTTGGGTTTATT-3’; mFoxp3_assay2_LR: 5’-ATAACTATATAATACATCAATACATTCTCA-3’; mFoxp3_ assay2_S2: 5’-GTTTTTTTTTTTTTTTTTTTGTTG-3’; mFoxp3_assay2_S3: 5’-GGTTGTGATAATAGGGTTTAGATGTAG-3’; and mFoxp3_assay2_S4: 5’-GTTTTTAAGAAATAGTTAAATAGG-3’ (15).

### Histopathological Analysis

Tissues were sectioned, fixed in 4% formaldehyde solution, embedded in paraffin, and stained with H&E. The sections were analyzed by light microscopy from Fisher Scientific Moticam at 20X magnification at room temperature. The images were acquired by using the software Motic Images Plus 2.0 (15).

### Quantitative Real-time RT-PCR Analysis

Total RNA was extracted with RNeasy mini kit from Qiagen. Isolated RNA was converted to cDNA by using High-Capacity cDNA Reverse Transcription Kit (Applied Biosystems, 4368814). Real-time RT-PCR was performed with Platinum SYBR Green qPCR SuperMix-UDG with ROX (Invitrogen, 11-744-500) and measured on StepOnePlus Real-Time PCR System (Applied Biosystems, 4376600). Data were normalized to 18S rRNA.

Primer sequences are as follows:

1. CAI: Forward 5’ GCGTTTTGATGAAGGTTGGT 3’ Reverse 5’ TCACCCAGGTCACACTTTCA 3’
2. DNMT3a: Forward 5’ ACTTGGAGAAGCGGAGTGAA 3’ Reverse 5’ GGATTCGATGTTGGTCTGCT 3’
3. TET1: Forward 5’ ATCATTCCAGACCGCAAGAC 3’ Reverse 5’ AATCCATGCAACAGGTGACA 3’
4. 18S: Forward 5’ GTAACCCGTTGAACCCCATT 3’ Reverse 5’ CCATCCAATCGGTAGTAGCG 3’

### RNA-seq

RNA was isolated from Treg cells with the RNeasy Mini Kit (Qiagen). The 75bp single-end RNA-seq was performed by the Genomics, Epigenomics, and Sequencing Core at the University of Cincinnati College of Medicine. Sequence reads were aligned to the reference mouse genome (mm10) using the TopHat aligner. Reads aligning to each known transcript were counted and all follow up analyses were performed using AltAnalyze software (25).

### Chromatin Immunoprecipitation-quantitative Real-time PCR (ChIP-qPCR)

Treg cells were fixed with 1% formaldehyde at room temperature for 10 min. Formaldehyde was quenched by addition of 0.125 M glycine and incubation for another 5 min. The cells were lysed in 1 ml buffer 1 (50 mM HEPES (pH 7.5), 140 mM NaCl, 1 mM EDTA, 0.5% NP-40, 0.25% Triton X-100, and 10% glycerol) supplemented with protease inhibitor cocktail (Roche, 36363600) for 10 min. Nuclei were collected and resuspended in 1 ml buffer 2 (10 mM Tris (pH 8.0), 200 mM NaCl, 0.5 mM EGTA and 1 mM EDTA), and incubated at 4 °C for 10 min. Pelleted nuclei were resuspended and sonicated for 10 min in 1 ml sonication buffer (100 mM NaCl, 50 mM Tris (pH 8), 5 mM EDTA pH 8, and 0.5% SDS) in a Covaris sonicator. After removal of an input control (whole cell lysate), chromatin was incubated with Anti-GATA3 antibody (Invitrogen, MA1-028, Clone ID: 1A12-1D9, RRID:AB_2536713) or normal mouse IgG (Santa Cruz, sc-2025, RRID:AB_737182) at 4 °C overnight, and pre-blocked Dynabeads® Protein G for Immunoprecipitation (Thermo Fisher Scientific, 10003D) were added to the samples. After incubation at 4 °C for 2 hr, the beads were washed with wash buffer 1 (150 mM NaCl, 20 mM Tris (pH 8), 5 mM EDTA (pH 8), 1% Triton X-100 and 0.2% SDS), Buffer 2 (0.1% deoxycholic acid, 1 mM EDTA (pH 8), 50 mM HEPES (pH 7.5), 1% Triton X-100, 500 mM NaCl), LiCl buffer (10 mM Tris (pH 8), 0.5% deoxycholic acid, 1 mM EDTA (pH 8), 250 mM LiCl, 0.5% NP-40), and Tris-EDTA (TE). DNA was eluted in elution buffer (0.1% SDS) and crosslinking was reversed by incubation at 65 °C for 10 hrs (36). After RNase A and proteinase K treatment, the precipitated chromatin DNA was purified by using a PCR purification kit (Qiagen, 28104) and then subjected to qPCR with the TaqMan™ Universal PCR Master Mix (Thermo Fisher Scientific, 4304437). Primers targeting the CAI locus containing GATA3 binding site are: Forward 5’ GTTGTCAGTTGCCTGGCATC 3’; Reverse: 5’ GACAGTGGTAGTGGCTGCAC 3’.

### Serum CASIN Detection

Blood was collected 7 hrs after the first CASIN injection and 14 hrs after the second CASIN injection into tumor-bearing mice. Serum was isolated using serum separator tubes. All serum samples were analyzed with Waters Quattro Micro UPLC system coupled to electrospray tandem mass spectrometry (ESIMS/MS). The separation was conducted on reverse phase Acquity® UPLC BEH C18 columns (100 × 0.1 mm. 1.7 μm) (Waters Crop. 186007488). The mobile phase was composed of two solvents: solvent A consisted of 50 ml/l acetonitrile and 950 ml/l water containing 2 mM ammonium acetate and 0.1% formic acid (v/v); Solvent B consisted of acetonitrile. The gradient conditions were used as described below with a flow rate of 0.1 ml/min. At time zero 85% A to 15% B, at 7 min 0% A to 100% B, keep this condition for 2 min and then change to 85% A to 15% B and keep for 3 min. The total run time was 15 min. 10 μl of sample was injected on column for analysis. Optimal MS signal for the ion of CASIN was achieved in negative ion mode using the following instrument settings: capillary voltage 3.0 kV; cone voltage 45 V, extractor voltage 2 V; RF lens voltage 0.1 V; entrance −1; exit 0; source temperature 120°C; desolvation temperature 400 °C; desolvation gas flow 600 l/hr; inter-channel delay 0.02 sec; inter-scan time 0.05 sec; dwell time 0.2 sec; helium was used as the collision gas and gas cell parani was about 3.6e-3. Optimal collision energy for CASIN was 30 V. CASIN was detected and quantified by monitoring the MRM transition ion m/z 305/244 under negative ESI mode. Data were acquired and processed with Masslynx 4.1 software (Waters Crop.) (17).

### DSS-induced Colitis

Colitis was induced in 8–10 week old mice by giving the mice 2.2% DSS (M.W. 36,000–50,000) in drinking water for 5 days followed by normal water (15). Mouse body weight was measured daily. The mice were sacrificed after 8 days and the colon was harvested for flow cytometry analysis.

### Immunoblotting

Whole-cell lysates were prepared and separated by 10% SDS-PAGE. Expression or activation (phosphorylation) of PAK1 and WASP was probed using the corresponding antibodies against total (Thermo Fisher Scientific, PA5-29224, RRID: AB_2546700) or phosphorylated (Thermo Fisher Scientific, PA5-37677, RRID: AB_2554285) PAK1 and total (Thermo Fisher Scientific, PA5-80224, RRID: AB_2747338) or phosphorylated (Thermo Fisher Scientific, PA5-38346, RRID:AB_2554947) WASP.

### Statistical Analysis

Tumor growth and body weight loss in DSS-induced mouse model of colitis were analyzed by two-way ANOVA. The rest of the statistics were performed with two-tailed Student t test. Data were expressed as mean ± SD. p < 0.05 was considered significant.

## Data Availability

Raw and processed data have been deposited in the National Center for Biotechnology Information GEO (accession no. GSE181779).

## Acknowledgements

This work was supported by grants from the National Institutes of Health (R01GM108661 to FG, R56 HL141499 to FG, R01CA234038 to FG and YZ, and R50CA211404 to MW), the National Center for Advancing Translational Sciences of the National Institutes of Health under Award Number 2UL1TR001425-05A1, and the Cincinnati Children’s Hospital Medical Center (RIP funding to FG).

We thank Veda Yadagiri and Hong Ji at the Cincinnati Children’s Hospital Research Foundation Pyrosequencing Core for analysis of methylation of Foxp3 gene and for writing the related methods.

## References

1. O.J. Finn, A Believer’s Overview of Cancer Immunosurveillance and Immunotherapy. J. Immunol. 200, 385–391 (2018).

2. A. Ribas, J.D. Wolchok, Cancer immunotherapy using checkpoint blockade. Science 359, 1350–1355 (2018).

3. P Sharma, S. Hu-Lieskovan, J.A. Wargo, A. Ribas, Primary, Adaptive, and Acquired Resistance to Cancer Immunotherapy. Cell 168, 707–723 (2017).

4. N. Ohkura, Y. Kitagawa, S. Sakaguchi, Development and maintenance of regulatory T cells. Immunity 38, 414–423 (2013).

5. E. V. Dang et al., Control of T(H)17/T(reg) balance by hypoxia-inducible factor 1. Cell 146, 772–784 (2011).

6. J. Gomez-Rodriguez et al., Itk-mediated integration of T cell receptor and cytokine signaling regulates the balance between Th17 and regulatory T cells. J. Exp. Med. 211, 529–543 (2014).

7. M. Liu, F. Guo, Recent updates on cancer immunotherapy. Precis. Clin. Med. 1, 65–74 (2018).

8. H. Nishikawa, S. Sakaguchi, Regulatory T cells in cancer immunotherapy. Curr. Opin. Immunol. 27, 1–7 (2014).

9. K. Shitara, H. Nhshikawa, Regulatory T cells: a potential target in cancer immunotherapy. Ann. N.Y. Acad. Sci. 1417, 104–115 (2018).

10. W.L. Byrne, K.H. Mills, J.A. Lederer, G.C. O’Sullivan, Targeting regulatory T cells in cancer. Cancer Res. 71, 6915–6920 (2011).

11. A. Colamatteo et al., Molecular Mechanisms Controlling Foxp3 Expression in Health and Autoimmunity: From Epigenetic to Post-translational Regulation. Front. Immunol. 10, 3136 (2019).

12. J. T. Cortez et al., CRISPR screen in regulatory T cells reveals modulators of Foxp3. Nature 582, 416–420 (2020).

13. R. Takahashi et al., SOCS1 is essential for regulatory T cell functions by preventing loss of Foxp3 expression as well as IFN-{gamma} and IL-17A production. J. Exp. Med. 208, 2055–2067 (2011).

14. J. Melendez, M. Grogg, Y. Zheng, Signaling role of Cdc42 in regulating mammalian physiology. J. Biol. Chem. 286, 2375–2381 (2011).

15. K.W. Kalim et al., Reciprocal regulation of glycolysis-driven Th17 pathogenicity and Treg stability by Cdc42. J. Immunol. 200, 2313–2326 (2018).

16. S. Parkkila, Significance of pH regulation and carbonic anhydrases in tumour progression and implications for diagnostic and therapeutic approaches. BJU. Int. 101, 16–21 (2008).

17. W. Liu et al., Rational identification of a Cdc42 inhibitor presents a new regimen for long-term hematopoietic stem cell mobilization. Leukemia 33, 749–761 (2019).

18. X. Li, Y. Liang, M. LeBlanc, C. Benner, Y. Zheng, Function of a Foxp3 cis-element in protecting regulatory T cell identity. Cell 158, 734–748 (2014).

19. G. Wei et al., Genome-wide analyses of transcription factor GATA3-mediated gene regulation in distinct T cell types. Immunity 35, 299–311 (2011).

20. Y. Wang, M.A. Su, Y.Y. Wan, An essential role of the transcription factor GATA-3 for the function of regulatory T cells. Immunity 35, 337–348 (2011).

21. S. Nomura et al., Pyrrothiogatain acts as an inhibitor of GATA family proteins and inhibits Th2 cell differentiation in vitro. Sci. Rep. 9, 17335 (2019).

22. F. Guo et al., Coordination of IL-7 receptor and T-cell receptor signaling by cell-division cycle 42 in T-cell homeostasis. Proc. Natl. Acad. Sci. U. S. A. 107, 18505–18510 (2010).

23. W.S. Lexmond et al., FOXP3+ Tregs require WASP to restrain Th2-mediated food allergy. J. Clin. Invest. 126, 4030–4044 (2016).

24. H. Kim, H. Falet, K.M. Hoffmeister, J.H. Hartwig, Wiskott-Aldrich Syndrome Protein (WASp) Controls the Delivery of Platelet Transforming Growth Factor-β-1. J. Biol. Chem. 288, 34352–34363 (2013).

25. J. Q. Yang et al., Rational targeting Cdc42 restrains Th2 cell differentiation and prevents allergic airway inflammation. Clin. Exp. Allergy 49, 92–107 (2019).

26. S. Sadhukhan, K. Sarkar, M. Taylor, F. Candotti, Y.M. Vyas, Nuclear role of WASp in gene transcription is uncoupled from its ARP2/3-dependent cytoplasmic role in actin polymerization. J. Immunol. 193, 150–160 (2014).

27. D. Brown, C.A. Wagner, Molecular Mechanisms of Acid-Base Sensing by the Kidney. J. Am. Soc. Nephrol. 23, 774–780 (2012).

28. R. Sakamori et al., CDC42 Inhibition suppresses progression of incipient intestinal tumors. Cancer Res. 74, 5480–5492 (2014).

29. L. E. Arias-Romero, J. Chernoff, Targeting Cdc42 in cancer. Expert Opin. Ther. Targets 17, 1263–1273 (2013).

30. A. P. Porter, A. Papaioannou, A. Malliri, Deregulation of Rho GTPases in cancer. Small GTPases 7, 123–138 (2016).

31. A. Czuchra et al., Cdc42 is not essential for filopodium formation, directed migration, cell polarization, and mitosis in fibroblastoid cells. Mol. Biol. Cell 16, 4473–4484 (2005).

32. F. Guo, C.S. Velu, H.L. Grimes, Y. Zheng, Rho GTPase Cdc42 is essential for B-lymphocyte development and activation. Blood 114, 2909–2916 (2009).

33. L. Yang et al., Rho GTPase Cdc42 coordinates hematopoietic stem cell quiescence and niche interaction in the bone marrow. Proc. Natl. Acad. Sci. U. S. A. 104, 5091–5096 (2007).

34. L. Yang et al., Cdc42 critically regulates the balance between myelopoiesis and erythropoiesis. Blood 110, 3853–3861 (2007).

35. M. Wunderlich et al., Improved multilineage human hematopoietic reconstitution and function in NSGS mice. PLoS One 13, e0209034 (2018).

36. C. Zhao et al., Dual regulatory switch through interactions of Tcf7l2/Tcf4 with stage-specific partners propels oligodendroglial maturation. Nat. Commun. 7, 10883 (2016).

