## Supplemental figures 1-11 for "Targeting of Cdc42 GTPase in regulatory T cells unleashes anti-tumor T cell immunity"

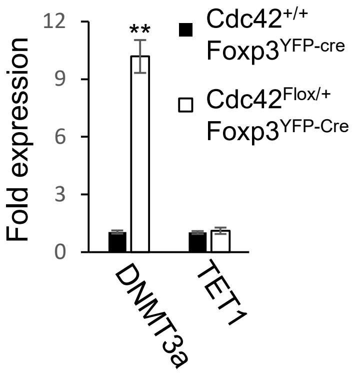


**Fig. S1.** *Cdc42* heterozygosity in Treg cells increases the expression of DNA methyltransferase DNMT3a but does not affect the expression of DNA demethylase TET1. The expression of DNMT3A and TET1 in $Cdc42^{+/+}Foxp3^{YFP\text{-}Cre}$ and $Cdc42^{Flox/+}Foxp3^{YFP\text{-}Cre}$ Treg cells was analyzed by quantitative real-time RT-PCR. Error bars indicate SD of triplicates. Data are from one experiment with four mice pooled. $^{**}p < 0.01$.

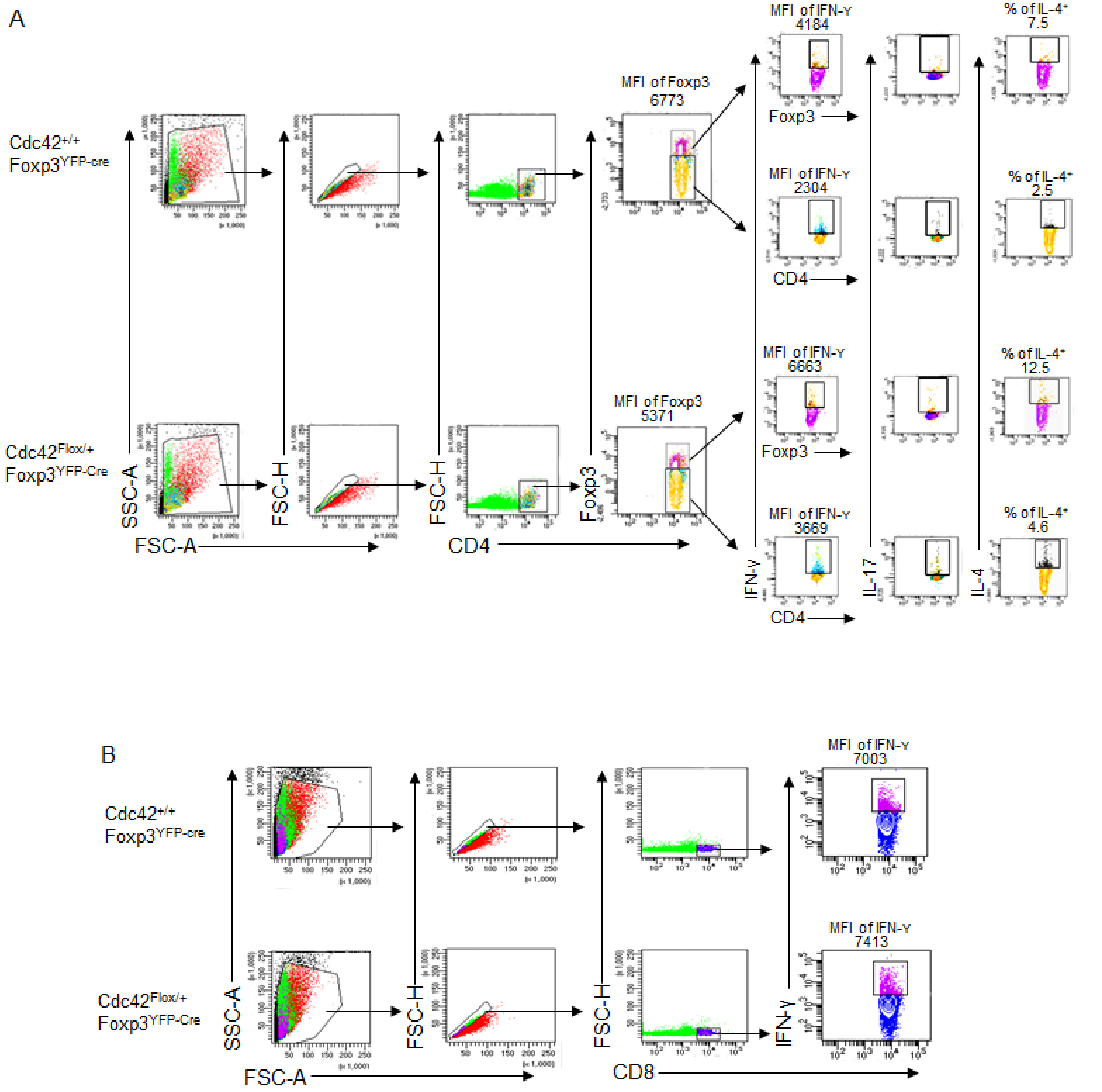


**Fig. S2.** Gating strategy and representative flow cytogram of Treg cells and CD4$^{+}$ and CD8$^{+}$ effector T cells expressing IFN-γ, IL-17 or IL-4. MFI: Mean fluorescence intensity.

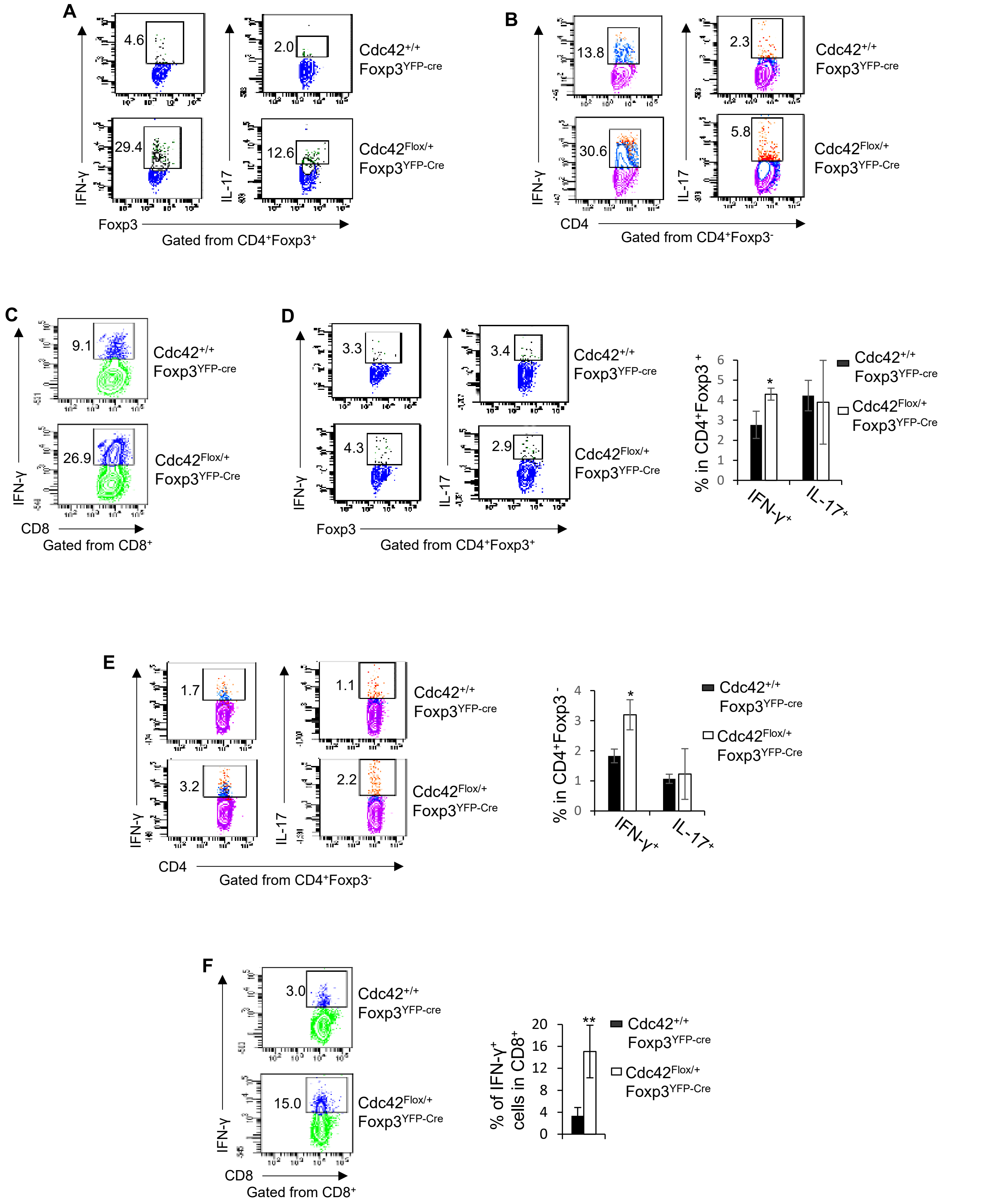


**Fig. S3.** *Cdc42* heterozygosity induces plasticity in both tumor-infiltrating and splenic Treg cells in tumor-bearing mice. (*A-C*) Representative flow cytogram of tumor-infiltrating Treg cells (*A*) and CD4$^{+}$ (*B*) and CD8$^{+}$ (*C*) effector

T cells expressing IFN-γ or IL-17. The numbers indicate percentages of tumor-infiltrating Treg cells and $CD4^+$ and $CD8^+$ effector T cells expressing IFN-γ or IL-17. (*D*-*F*) Left, representative flow cytogram of splenic Treg cells (*D*) and $CD4^+$ (*E*) and $CD8^+$ (*F*) effector T cells expressing IFN-γ or IL-17. The numbers indicate percentages of splenic Treg cells and $CD4^+$ and $CD8^+$ effector T cells expressing IFN-γ or IL-17. Right, average percentages of splenic Treg cells and $CD4^+$ and $CD8^+$ effector T cells expressing IFN-γ or IL-17. Error bars indicate SD of 6 mice. Data are representative of two independent experiments. $*p < 0.05$; $**p < 0.01$.

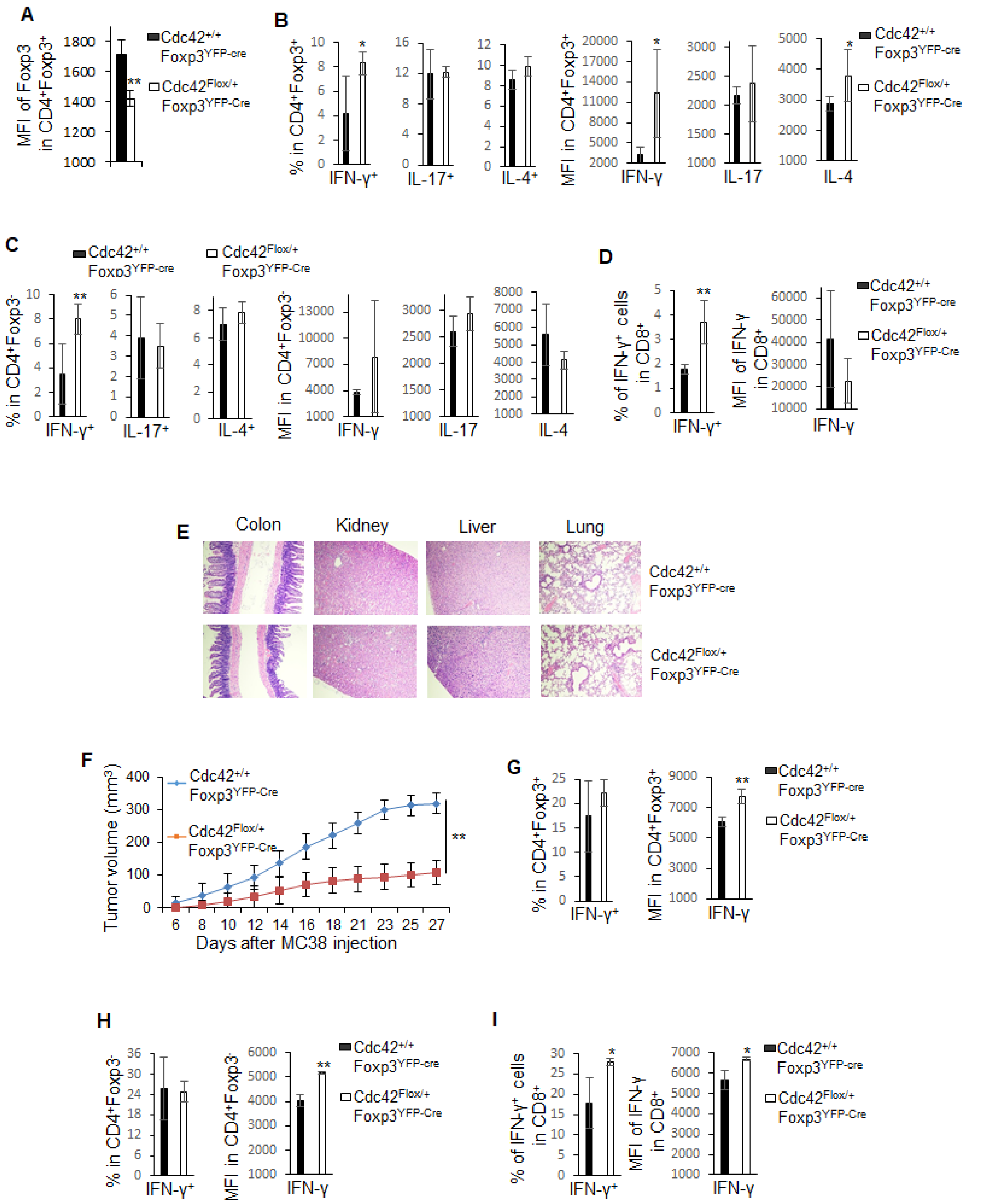


**Fig. S4.** *Cdc42* heterozygosity in Treg cells induces Treg cell plasticity and anti-tumor T cell immunity without causing autoimmunity in aged (12-18 month old) mice. (*A*) Flow cytometry analysis of the expression (MFI) of

Foxp3 in Treg cells. (*B-D*) Flow cytometry analysis of the expression (percentages and MFI) of IFN-γ, IL-17 and/or IL-4 in Treg cells (*B*), $CD4^+$ T cells (*C*) and $CD8^+$ T cells (*D*). (*E*) H&E staining of the indicated organs. (*F*) Tumor growth of MC38 mouse colon cancer cells. (*G-I*) Flow cytometry analysis of the expression (percentages and MFI) of IFN-γ in MC38 tumor-infiltrating Treg ($CD4^+Foxp3^+$) (*G*), $CD4^+$ effector T ($CD4^+Foxp3^-$) (*H*) and $CD8^+$ effector T cells (*I*). Error bars indicate SD of 5 mice. $^*p < 0.05$; $^{**}p < 0.01$. MFI: Mean fluorescence intensity.

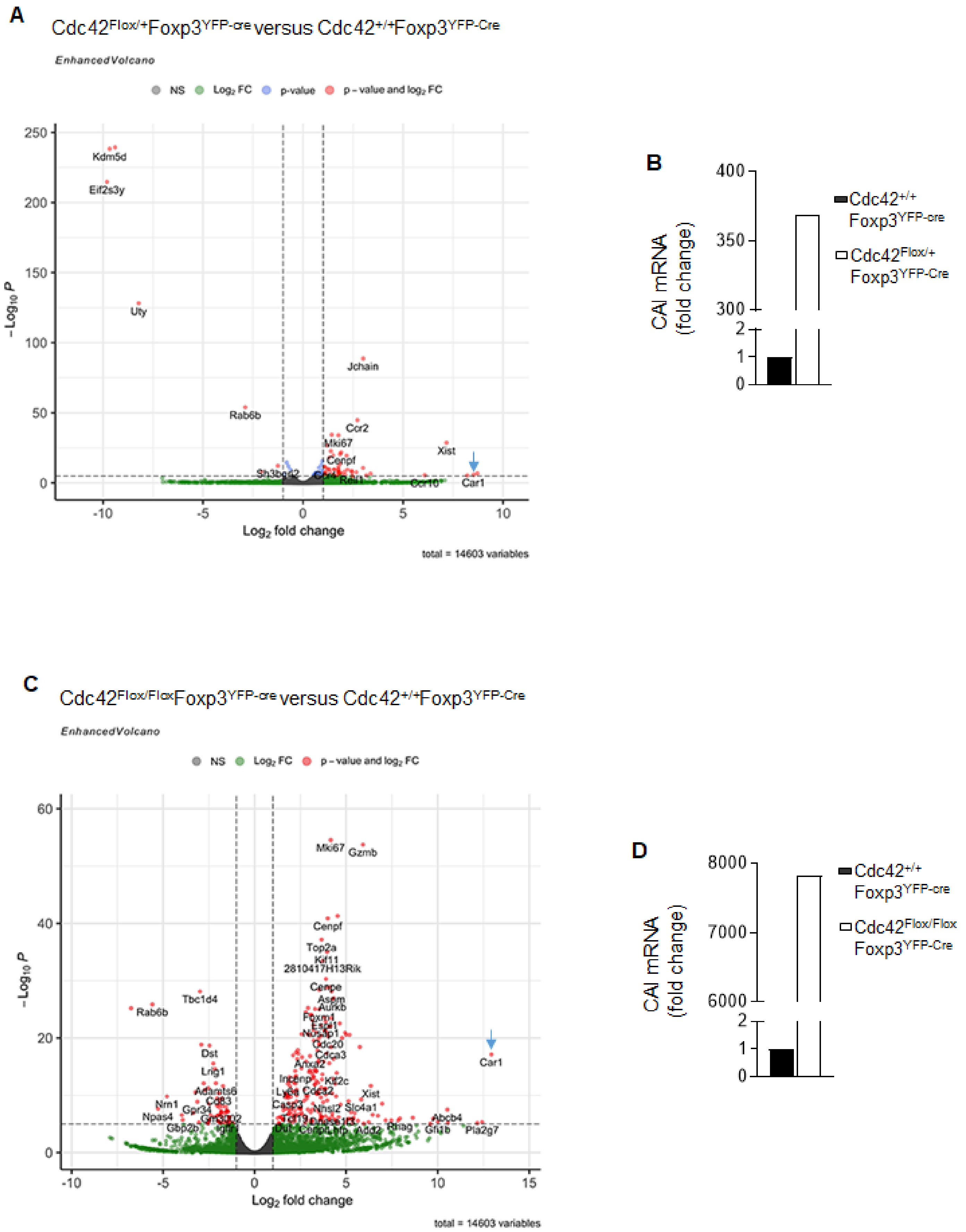

A
Cdc42Flox/+Foxp3YFP-cre versus Cdc42+/+Foxp3YFP-Cre
EnhancedVolcano
NS
Log2 FC
p-value
p – value and log2 FC
–Log10 P
Log2 fold change
total = 14603 variables
Kdm5d
Eif2s3y
Uty
Jchain
Rab6b
Ccr2
Mki67
Xist
Cenpf
Sh3bgrl2
Car1
Ccr10
B
CAI mRNA
(fold change)
Cdc42+/+
Foxp3YFP-cre
Cdc42Flox/+
Foxp3YFP-Cre
C
Cdc42Flox/FloxFoxp3YFP-cre versus Cdc42+/+Foxp3YFP-Cre
EnhancedVolcano
NS
Log2 FC
p – value and log2 FC
–Log10 P
Log2 fold change
total = 14603 variables
Mki67
Gzmb
Cenpf
Top2a
Kif11
2810417H13Rik
Cenpe
Aspm
Aurkb
Tbc1d4
Rab6b
Car1
Dst
Lrig1
Adamts6
Nrn1
Npas4
Gpr34
Gbp2b
Arhgap11a
Kif2c
Xist
Slo4a1
Abcb4
Pla2g7
Rhag
Add2
D
CAI mRNA
(fold change)
Cdc42+/+
Foxp3YFP-cre
Cdc42Flox/Flox
Foxp3YFP-Cre

**Fig. S5.** Global gene expression profiling of Treg cells bearing heterozygous or homozygous *Cdc42* deletion. (*A*) Volcano plot of gene expression changes in *Cdc42*$^{Flox/+}$*Foxp3*$^{YFP\text{-}cre}$ Treg cells in comparison with *Cdc42*$^{+/+}$*Foxp3*$^{YFP\text{-}Cre}$ Treg cells. Blue arrow points to *Car1* gene that encodes CAI. (*B*) Fold change of CAI in *Cdc42*$^{Flox/+}$*Foxp3*$^{YFP\text{-}cre}$ Treg cells, compared to *Cdc42*$^{+/+}$*Foxp3*$^{YFP\text{-}Cre}$ Treg cells. (*C*) Volcano plot of gene expression changes in *Cdc42*$^{Flox/Flox}$*Foxp3*$^{YFP\text{-}cre}$ Treg cells in comparison with *Cdc42*$^{+/+}$*Foxp3*$^{YFP\text{-}Cre}$ *Treg* cells. Blue arrow points to Car1 gene. (*D*) Fold change of CAI in *Cdc42*$^{Flox/Flox}$*Foxp3*$^{YFP\text{-}cre}$ Treg cells, compared to *Cdc42*$^{+/+}$*Foxp3*$^{YFP\text{-}Cre}$ Treg cells.

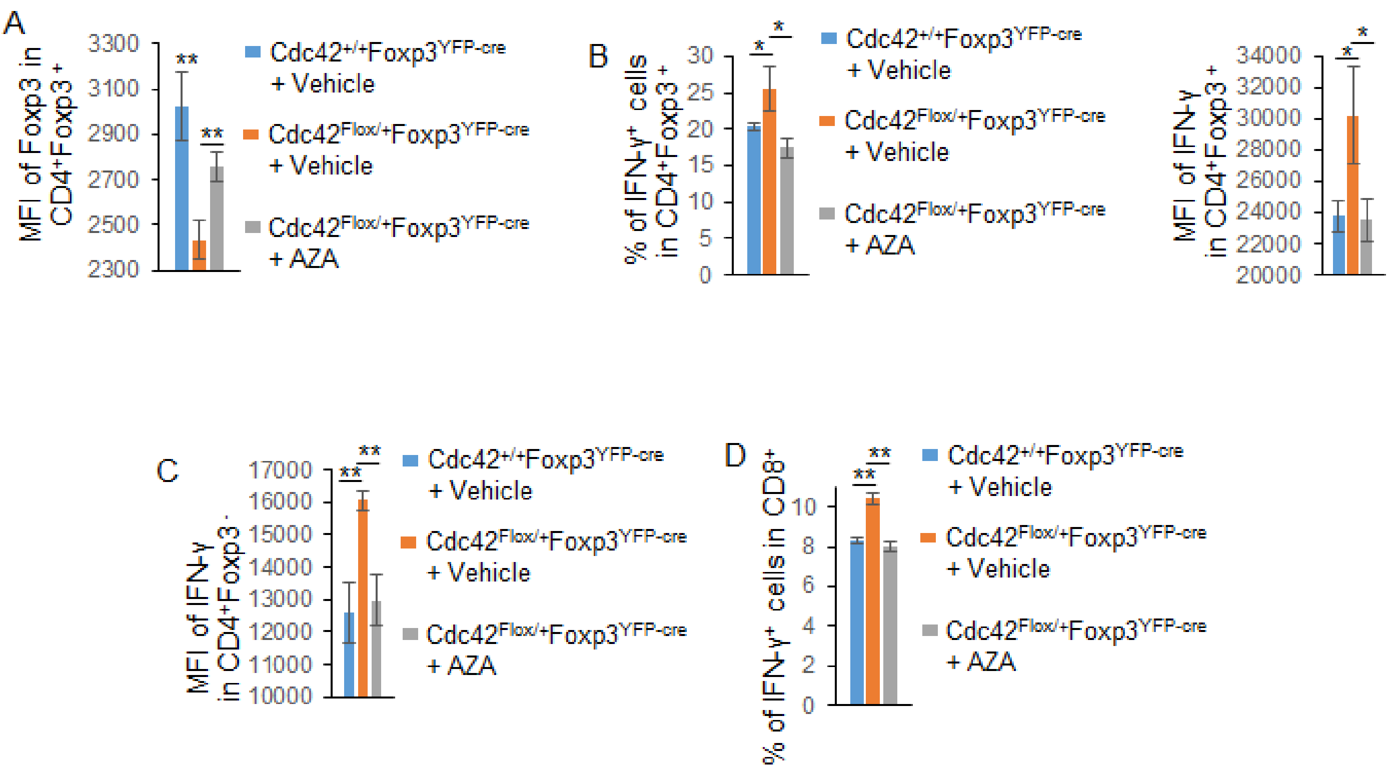


**Fig. S6.** Inhibition of CAI in *Cdc42* haploinsufficient mice reduces plasticity of tumor-infiltrating Treg cells and inhibits tumor-infiltrating effector T cells. (*A*) Flow cytometry analysis of the expression of Foxp3 in MC38 tumor-infiltrating Treg cells from *Cdc42*$^{+/+}$*Foxp3*$^{YFP\text{-}Cre}$ and *Cdc42*$^{Flox/+}$*Foxp3*$^{YFP\text{-}Cre}$ mice treated with or without AZA. (*B-D*) Flow cytometry analysis of the expression of IFN-γ in MC38 tumor-infiltrating Treg cells (*B*), CD4$^{+}$ effector T cells (*C*) and CD8$^{+}$ effector T cells (*D*) from *Cdc42*$^{+/+}$*Foxp3*$^{YFP\text{-}Cre}$ and *Cdc42*$^{Flox/+}$*Foxp3*$^{YFP\text{-}Cre}$ mice treated with or without AZA. Error bars indicate SD of 4 mice. Data are representative of two independent experiments. $^{*}p < 0.05$; $^{**}p < 0.01$. AZA: acetazolamide. MFI: Mean fluorescence intensity.

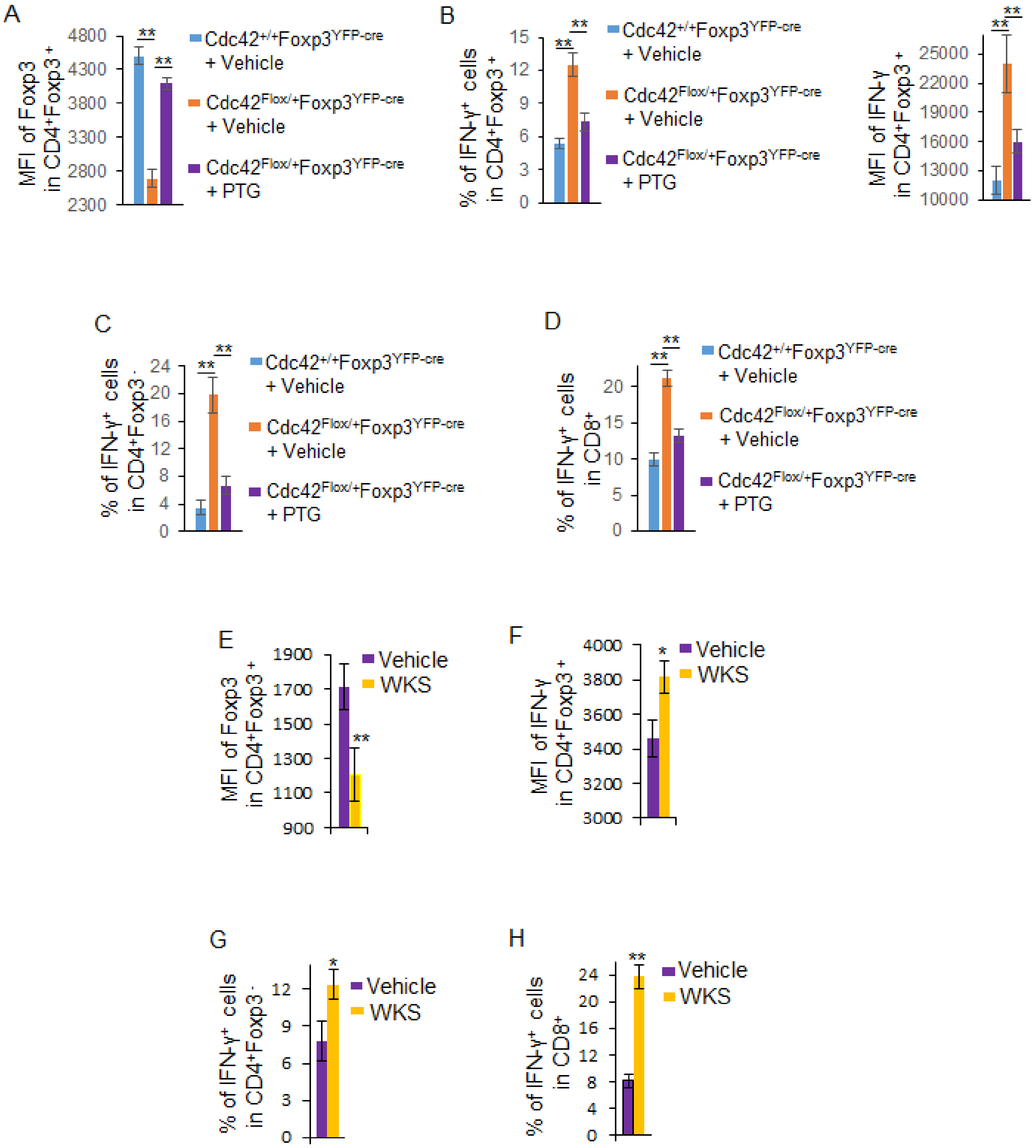


**Fig. S7.** Inhibition of GATA3 in *Cdc42* haploinsufficient mice reduces plasticity of tumor-infiltrating Treg cells and inhibits tumor-infiltrating effector T cells and inhibition of WASP mimics heterozygous loss of *Cdc42* in inducing tumor-infiltrating Treg cell plasticity and increasing tumor-infiltrating effector T cells. (*A*) Flow cytometry analysis of the expression of Foxp3 in MC38 tumor-infiltrating Treg cells from *Cdc42*$^{+/+}$*Foxp3*$^{YFP\text{-}Cre}$ and *Cdc42*$^{Flox/+}$*Foxp3*$^{YFP\text{-}Cre}$ mice treated with or without PTG. (*B-D*) Flow cytometry analysis of the expression of IFN-γ in MC38 tumor-infiltrating Treg cells (*B*), CD4$^{+}$ effector T cells (*C*) and CD8$^{+}$ effector T cells (*D*) from *Cdc42*$^{+/+}$*Foxp3*$^{YFP\text{-}Cre}$ and *Cdc42*$^{Flox/+}$*Foxp3*$^{YFP\text{-}Cre}$ mice treated with or without PTG. (*E*) Flow cytometry analysis of the expression of Foxp3 in MC38 tumor-infiltrating Treg cells from *Cdc42*$^{+/+}$*Foxp3*$^{YFP\text{-}Cre}$ mice treated with or without WKS. (*F-H*) Flow cytometry analysis of the expression of IFN-γ in MC38 tumor-infiltrating Treg cells (*F*), CD4$^{+}$ effector T cells (*G*) and CD8$^{+}$ effector T cells (*H*) from *Cdc42*$^{+/+}$*Foxp3*$^{YFP\text{-}Cre}$ mice treated with or without WKS. Error bars indicate SD of 4 mice. Data are representative of two independent experiments. *p < 0.05; **p < 0.01. PTG: pyrrothiogatain. WKS: wiskostatin. MFI: Mean fluorescence intensity.

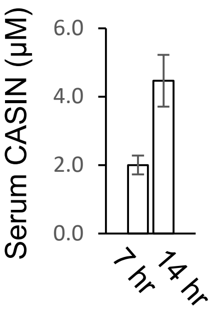


**Fig. S8.** Serum levels of CASIN. C57/BL6 mice were treated with CASIN starting upon MC38 tumor onset. Serum was collected 7 hr after the first CASIN (30 mg/Kg) injection and 14 hr after the second CASIN (30 mg/Kg) injection that was performed 9 hr after the first injection. Error bars indicate SD of 4 mice.

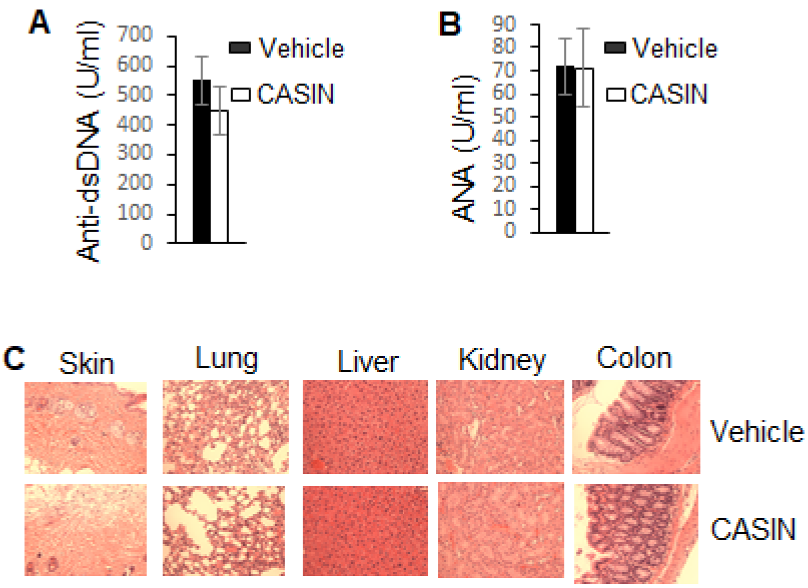


**Fig. S9.** Tumor-bearing mice therapeutically treated with CASIN do not show systemic autoimmune responses. (*A* and *B*) ELISA analysis of the concentrations of serum autoantibodies from C57/BL6 mice treated with or without CASIN starting upon MC38 tumor onset. (*C*) H&E staining of the indicated organs. (*A* and *B)* Error bars indicate SD of 6 mice. Anti-dsDNA: Anti-double stranded DNA. ANA: Antinuclear antibody.

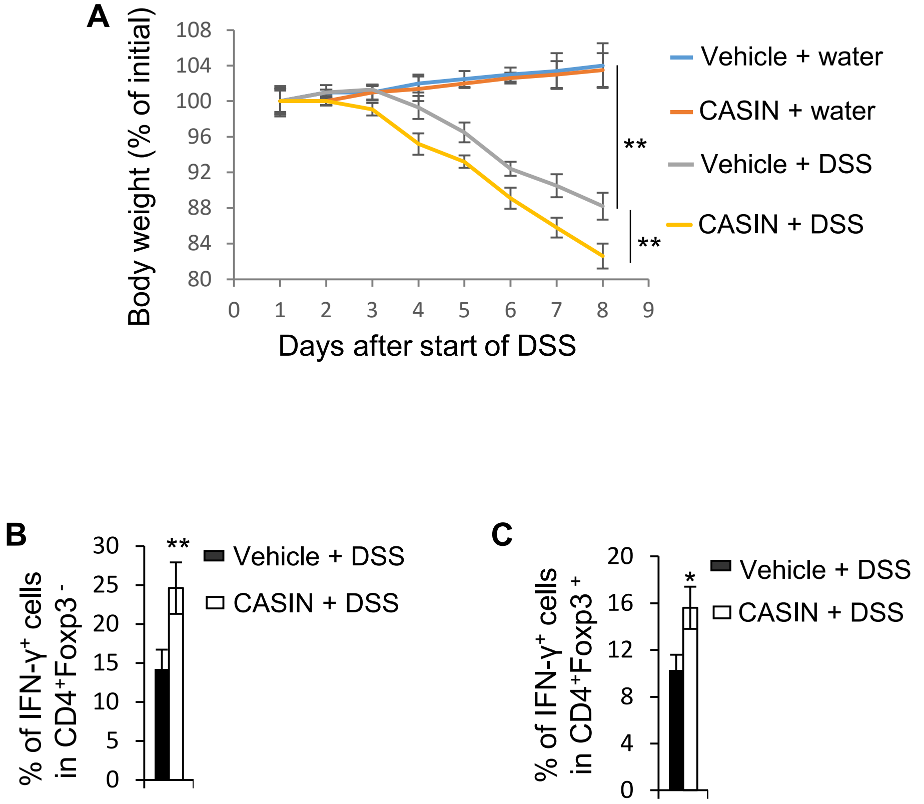


**Fig. S10.** CASIN increases disease severity of colitis. (*A*) Body weight loss of C57BL/6 mice treated with or without CASIN and/or DSS. The data are expressed as percentage of initial body weight. (*B* and *C*) Percentages of IFN-γ-producing colonic $CD4^+$ effector T cells (*B*) and Treg cells (*C*). Error bars indicate SD of 6 mice. $^*p < 0.05$; $^{**}p < 0.01$.

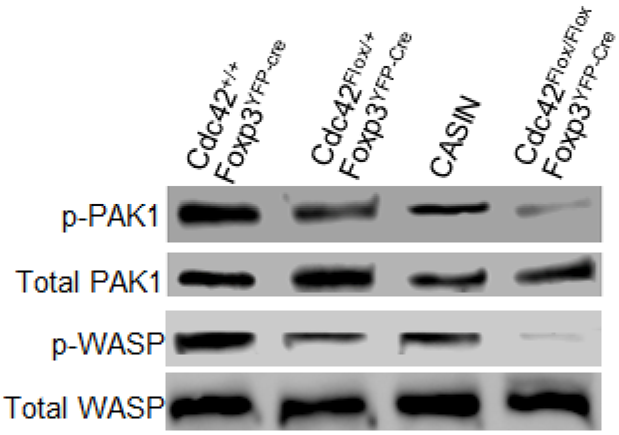


**Fig. S11.** Heterozygous *Cdc42* deletion and CASIN inhibit Cdc42 effector activation to a lesser extent than homozygous *Cdc42* deletion. Splenic Treg cells from *Cdc42*$^{+/+}$*Foxp3*$^{YFP\text{-}Cre}$, *Cdc42*$^{Flox/+}$*Foxp3*$^{YFP\text{-}Cre}$, and *Cdc42*$^{Flox/Flox}$*Foxp3*$^{YFP\text{-}Cre}$ mice were treated with or without CASIN (2 μM) for 3 days. Phosphorylated (p-) and total PAK1 and WASP were analyzed by Western blot.
